# The CHK-2 antagonizing phosphatase PPM-1.D regulates meiotic entry via catalytic and non-catalytic activities

**DOI:** 10.1101/2021.08.02.453806

**Authors:** Antoine Baudrimont, Dimitra Paouneskou, Ariz Mohammad, Raffael Lichtenberger, Joshua Blundon, Yumi Kim, Markus Hartl, Sebastian Falk, Tim Schedl, Verena Jantsch

## Abstract

The transition from the stem cell/progenitor fate to meiosis is mediated by several redundant post-transcriptional regulatory pathways in *C. elegans*. Interfering with all three branches causes tumorous germlines. SCF^PROM-1^ comprises one branch and mediates a scheduled degradation step at entry into meiosis. *prom-1* mutants show defects in timely initiation of events of meiotic prophase I, resulting in high rates of embryonic lethality. Here, we identify the phosphatase PPM-1.D/Wip1 as crucial substrate for PROM-1. We report that PPM-1.D antagonizes CHK-2 kinase, a key regulator for meiotic prophase initiation e.g., DNA double strand breaks, chromosome pairing and synaptonemal complex formation. We propose that PPM-1.D controls the amount of active CHK-2 by both catalytic and non-catalytic activities, where strikingly the non-catalytic regulation seems to be crucial at meiotic entry. PPM-1.D sequesters CHK-2 at the nuclear periphery and programmed SCF^PROM-1^ mediated degradation of PPM-1.D liberates the kinase and promotes meiotic entry.

## Introduction

The transition from the dividing stem/progenitor cell fate to meiosis is a key step in producing gametes (Hubbard and Schedl, 2019). In the germline this crucial differentiation step is governed by three parallel pathways involved in post-transcriptional gene regulation in *C. elegans*. These include the GLD-1, GLD-2 and SCF^PROM-1^ pathways that act by translational repression, polyA tail mediated translational activation and targeted protein degradation, respectively (Mohammad et al., 2018). The pathways operate redundantly, which means that only double mutants interfering with at least two pathway branches lead to over-proliferative germlines and failure in meiotic entry. Triple mutants affecting all three pathways produce highly tumorous germlines with little or no expression of meiotic markers (Mohammad et al., 2018). In the progenitor zone, where cells undergo mitotic cell cycling and pre-meiotic replication, the activities of the three pathways required for meiotic entry are downregulated by GLP-1/Notch signaling (Hansen et al., 2004; Mohammad et al., 2018).

The continuous replenishment of meiocytes through divisions in the progenitor zone displaces cells proximally at a rate of approximately one cell row/hour through the germline (Crittenden et al., 2006). After one round of meiotic S-phase, cells enter prophase of meiosis I (leptonema, zygonema, pachynema, diplonema, and diakinesis), which is organized as a spatio-temporal meiotic time course in the dissected gonads of *C. elegans* hermaphrodites (Hillers et al., 2017). The generation of gametes via meiosis requires two divisions. In meiosis I, parental homologous chromosomes (one from each parent) are separated and in meiosis II, each chromosome splits into its two sister chromatids.

The physical linkage between homologs aids their correct segregation. This linkage is a result of programmed induction of DNA double strand breaks (DSBs), pairwise alignment of the homologous chromosomes, which are organized in loops tethered to the meiotic chromosome axis, installation of the synaptonemal complex (SC) between the paired homologs and repair of the DSBs using a chromatid of the parental homolog via homologous recombination (Gerton and Hawley, 2005). A further highly conserved feature in prophase of meiosis is the chromosome end led movements, which promote the pairwise alignment of the homologous chromosomes and the installation of the SC between them (Link and Jantsch, 2019). These events must be coordinated to achieve normal disjunction at the meiotic divisions.

*prom-1* mutants show defects in timely and coordinated initiation of these events (Jantsch et al., 2007). The mutants have an extended meiotic entry zone, characterized by the presence of meiotic cohesion, chromosome axes and SC proteins as poly-complexes, indicating that the proteins are produced and await assembly onto chromosomes. Furthermore, despite apparent completion of meiotic S-phase, DSB induction and repair and all signs of the prophase chromosome movements are delayed. These pleiotropic defects result in a mix of univalent and bivalents, which leads to chromosome mis-segregation and high embryonic death (Jantsch et al., 2007).

In *C. elegans*, the DNA damage signaling kinase CHK-2 acts as a key regulator of prophase meiotic processes. *chk-2* mutants are defective in DSB induction, SC formation, chromosome movements and lack meiotic feedback control that permits bivalent formation (Castellano-Pozo et al., 2020; Kim et al., 2015; Link et al., 2018; MacQueen and Villeneuve, 2001; Penkner et al., 2009; Rosu et al., 2013; Sato et al., 2009; Stamper et al., 2013; Woglar and Jantsch, 2013). The nuclear envelope protein SUN-1, which is involved in the chromosome movements, is a prominent substrate of CHK-2 and phosphorylated SUN-1 serine 8 (SUN-1(S8Pi)) marks meiotic entry (Penkner et al., 2009) and is used as a marker for CHK-2 kinase activity throughout this study. Fundamentally different from the *prom-1* mutants, *chk-2* mutants show normal axes morphogenesis (Tang et al., 2010).

*prom-1* encodes an F-box protein homologous to human FBX047 (Jantsch et al., 2007; Simon-Kayser et al., 2005). Together with a cullin and an Rbx protein, PROM-1 is part of a multi-subunit E3 ubiquitin ligase complex (called SCF) (Nayak et al., 2002), which mediates recognition and binding of the E2 ubiquitin-conjugating enzyme to the substrate, which is consecutively targeted for degradation. We still do not have a comprehensive picture of which proteins need to be subjected to the programmed degradation step at the transition between the stem/progenitor cell fate and meiotic differentiation. Whereas the cyclin, CYE-1, has been identified as one of the targets of SCF^PROM-1^, *cye-1* inactivation failed to rescue the pronounced meiotic entry delay seen in *prom-1* mutant worms (Mohammad et al., 2018).

In this study, we report the identification of *ppm-1.D* as a potent suppressor of the embryonic lethality associated with the *prom-1* mutants. *prom-1* defects in meiotic entry are largely reversed and key meiotic processes of prophase I restored. *ppm-1.D* encodes a serine/threonine phosphatase in the PP2C family that is orthologous to human *PPM1D* (formally known as *WIP1*). We provide evidence that PPM-1.D acts as an antagonizing phosphatase to the meiotic regulator CHK-2, which it can keep inactive by a mere sequestration mechanism (non-catalytic activity) in the progenitor zone compartment. Nevertheless, PPM-1.D regulates meiotic entry via both catalytic and non-catalytic activities and therefore *ppm-1.D* null mutants display features of premature meiotic entry. Thus we present a yet undescribed role for the PPM-1.D phosphatase, besides its known involvement in the response to DNA damage in somatic cells (Le Guezennec and Bulavin, 2010). This study provides incentives to test whether human PPM1D is also a substrate for degradation by the human *prom-1* F-box protein homolog, FBX047, where mutations in the gene have been associated with renal carcinoma (Simon-Kayser et al., 2005). Furthermore, *PPM1D* is often found up-regulated in cancer cells (Le Guezennec and Bulavin, 2010).

## Results

### Identification of *ppm-1.D* as a *prom-1* target

*prom-1* encodes an F-box protein and is part of the SCF E3 ubiquitin ligase complex, which targets substrate proteins for degradation by the proteasome (Jantsch et al., 2007; Mohammad et al., 2018; Nayak et al., 2002) (Figure 1.A). We tagged PROM-1 at its carboxy-terminus and examined its expression levels throughout the *C. elegans* germline (see Table S1 for functionality of the tagged line). We co-stained PROM-1::HA with the cohesion regulator WAPL-1, which shows a characteristic nuclear staining in the progenitor zone with a pronounced drop at meiotic entry (Crawley et al., 2016) (Figure 1.B, left, cyan). Quantification of the normalized signal intensity of PROM-1 revealed that it started to rise ∼10 cells diameters (rows) from the distal tip of the germline and reached its maximum level ∼20 cell diameters from the distal tip (Figure 1.B). The peak is ∼20 fold above the base in the distal most germ cells and coincides with the end of the progenitor zone marked by WAPL-1 (Figure 1. B, right, cyan triangle). The increase in the levels of PROM-1 right at meiotic entry suggests the presence of targets for regulated degradation to promote entry into meiosis, consistent with the *prom-1* mutant phenotype with the characteristic extended meiotic entry zone (Jantsch et al., 2007).

**Figure 1.**
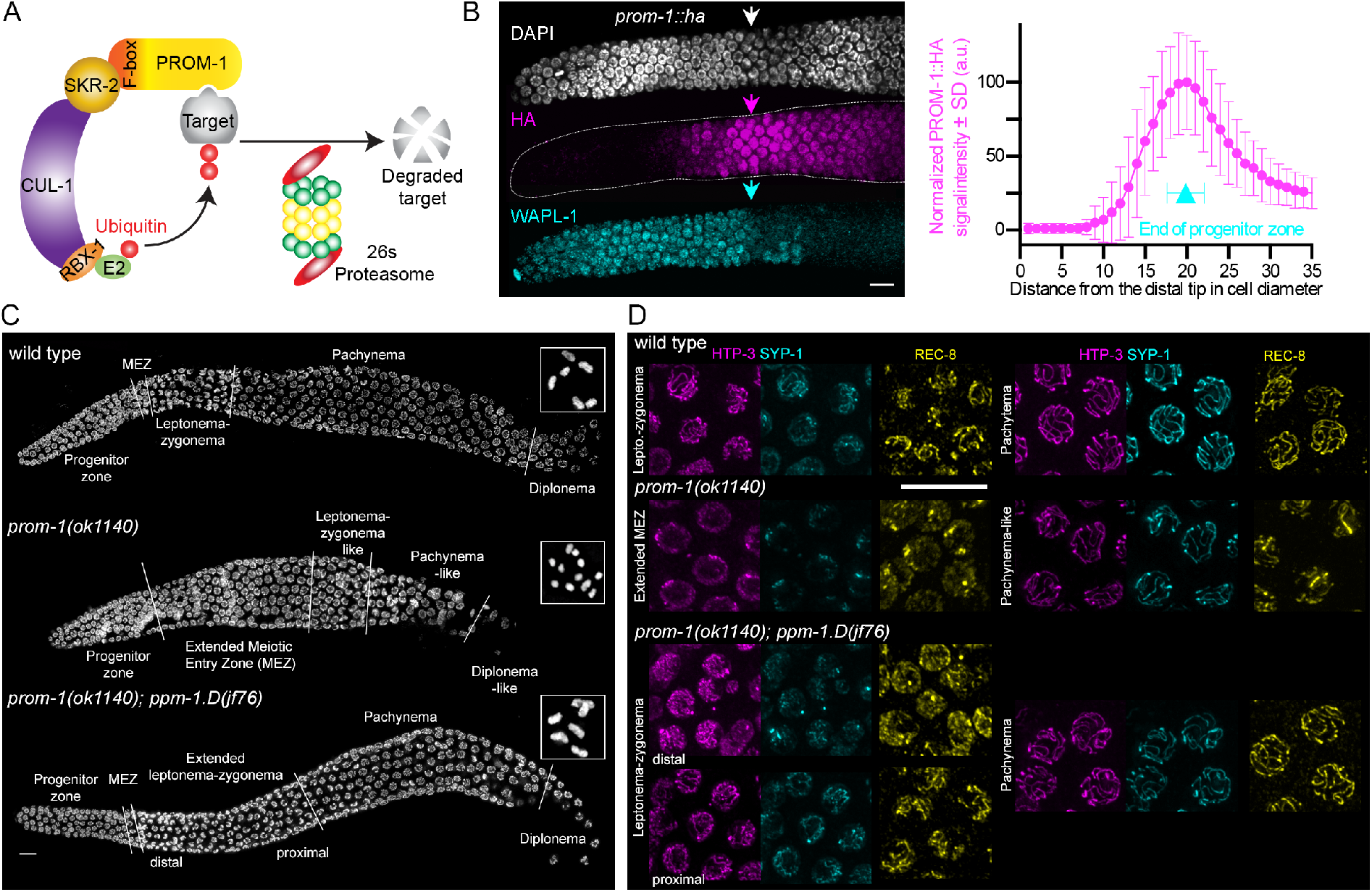
Loss of SCF^PROM-1^ activity at meiotic entry is rescued by mutating *ppm-1.D.* **A**. Schematics of the SCF^PROM-1^ complex. **B**. Left, Immunodetection of WAPL-1 (cyan) and PROM-1::HA (magenta) in the progenitor zone, at the distal end of the *C. elegans* germline. Arrows marks the entry into meiosis, which occurs at the leptonema-zygonema stage. Scale bar: 5 µm. Right, normalized levels of PROM-1::HA (magenta) throughout the progenitor cell zone, measured along the distance from the distal tip in cell diameter; the end of the progenitor zone (cyan) is marked. Error bars: SD. **C**. Gonads displaying prophase I for the mentioned genotypes. Scale bar: 10 µm. Boxed insets show representative diakinesis chromosomes. **D**. Insets showing staining for HTP-3 (magenta), SYP-1 (cyan), REC-8 (yellow) for the depicted zones. Scale bar: 10 µm.

To identify targets of SCF^PROM-1^, we conducted a suppressor screen in search of mutants that would rescue the low viability of *prom-1(ok1140)* (15 ± 10%, n = 7 hermaphrodites) (see materials and methods and Figure S1.A). We isolated the allele *jf76*, which mapped to the *ppm-1.D* gene. Combining *jf76* with *prom-1* leads to a significantly improved hatch rate of 79 ± 14% (n = 10 hermaphrodites) (Figure S1.B).

Further cytological examination of the double mutant *prom-1(ok1140); ppm-1.D(jf76)* revealed: 1) the timely restoration of the appearance of the leptonema-zygonema after the meiotic entry zone (MEZ, comprising the 2-3 nuclear cell rows in the wild type, where SC proteins have been expressed, but not yet loaded onto chromosomes (Jantsch et al., 2007)) contrasting the extended MEZ in *prom-1(ok1140)* (Figure 1.C), 2) the loading of the meiotic cohesion REC-8 and chromosome axial proteins (as shown for HTP-3 (Goodyer et al., 2008)), and extension of the SC (as shown for SYP-1 (MacQueen et al., 2002; Schild-Prufert et al., 2011)) (Figure 1.D). We noticed that in the double mutant the transition zone (comprising leptonema and zygonema) was prolonged and that HTP-3, SYP-1 and REC-8 persisted longer in aggregates than in the wild type. Nevertheless, at the proximal end of the transition zone the chromosome axes and the SC appeared fully decorated with the relevant markers (Figure 1.D) and 3) and the formation of six bivalents compared to the mixture of univalent and bivalents in the *prom-1(ok1140)* single mutant (Figure 1.C, insets). Consistent with the efficient formation of bivalents, pairing of homologous chromosomes and RAD-51 loading were restored to wild-type levels in the *prom-1(ok1140); ppm-1.D(jf76)* double mutant (Figure S1.C,D).

In summary, we showed that PROM-1 protein levels peak at meiotic entry and that the *ppm-1.D(jf76)* mutant can efficiently suppress the *prom-1* phenotype as evidenced by restoration of high hatch rates of embryos laid by the double mutant.

### PPM-1.D encodes a conserved PP2C phosphatase and protein abundance is regulated by the SCF^PROM-1^ complex

The *ppm-1.D* gene is conserved from *C. elegans* to human (Figure 2.A) and is known for its involvement in the DNA damage response in mammals (Goloudina et al., 2016). PPM1.D is a chromatin-bound phosphatase targeting γH2AX, ATM, CHK1, CHK2, MDM2, and p53 and reverses effects of ATM-dependent mitotic cell cycle arrest triggered by DNA damage. In animal cells, the amount of the chromatin-bound PPM1D/WIP1-ATM complex regulates the duration of cell cycle arrest after DNA damage induction (Jaiswal et al., 2017).

**Figure 2.**
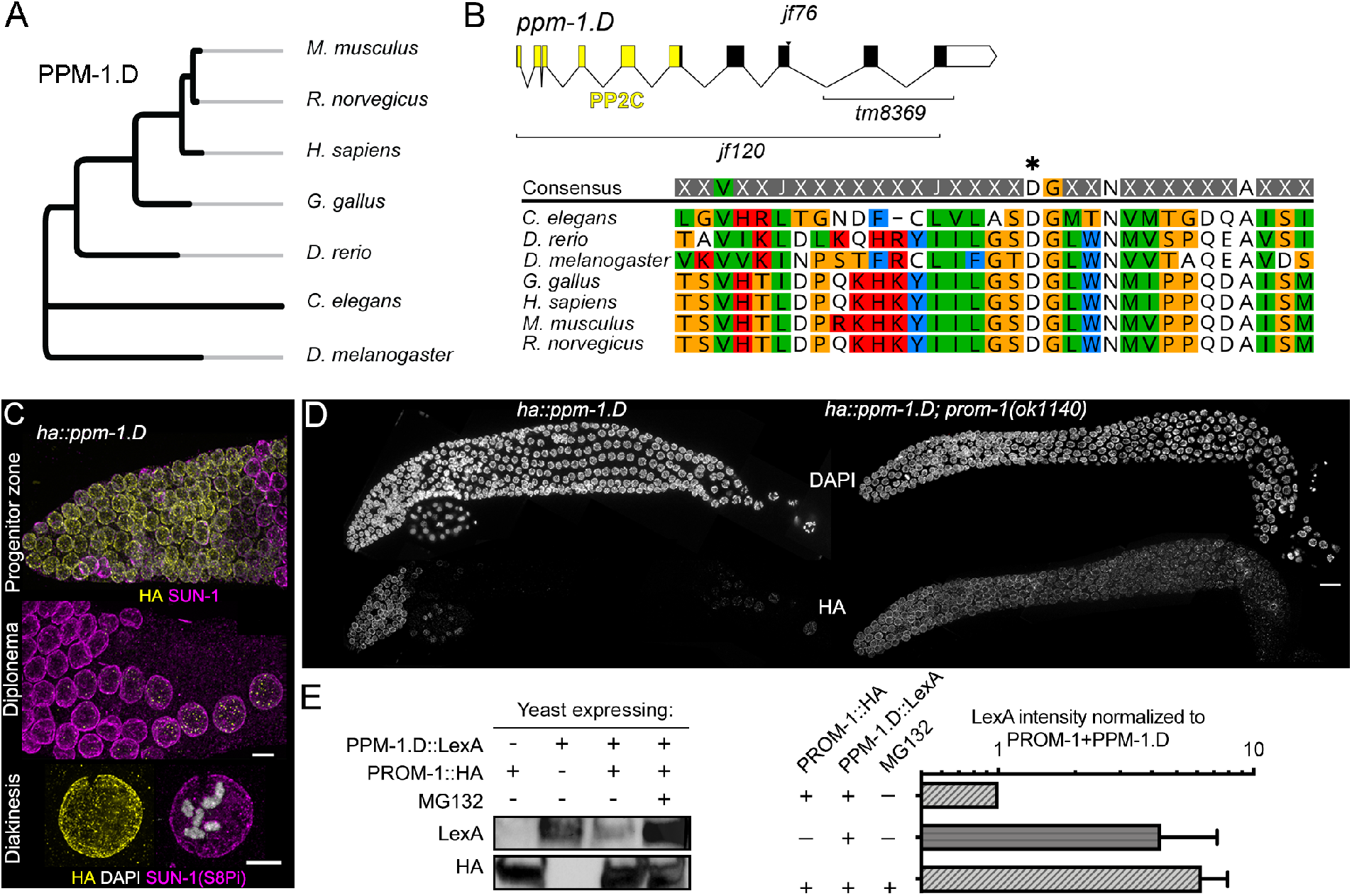
PPM-1.D is a conserved PP2C phosphatase and expression is controlled by SCF^PROM-1.^ **A**. Phylogenetic tree of PPM-1.D. **B**. Gene structure of *ppm-.1D* with domains, exons/introns and alleles depicted (top), and alignment of PPM-1.D protein sequences (bottom) (amino acid range: 498 – 530) for the mentioned organisms highlighting the conservation of the PP2C domain. Asterisk marks the conserved aspartic acid necessary for phosphatase activity. **C**. Immuno-detection of PPM-1.D::HA (yellow) and SUN-1 (magenta) in the progenitor zone (top), diplonema (middle) and diakinesis (bottom). Scale bar: 5 µm. **D**. Dissected gonads stained for DAPI (top) and PPM-1.D::HA in wild type (left) and *prom-1* mutants (right). Scale bar: 10 µm. **E**. Left, Western blot with TCA precipitated proteins from yeast expressing PPM-1.D::LexA, PROM-1::HA in absence or presence of the proteasome inhibitor. Right, quantification of PPM-1.D::LexA in the western blots (n=2) normalized to the level of PPM-1.D::LexA when both PROM-1 and PPM-1.D are expressed.

*C. elegans* PPM-1.D has a phosphatase type 2C domain (PP2C) (Figure 2.B) classifying it as a member of the corresponding phosphatases family (Bork et al., 1996). The allele *jf76,* which suppresses the high level of embryonic death in the *prom-1* mutant, bears a G to C transversion that abolishes splicing and leads to a premature stop codon. This leads to the loss of the last two exons similar to the *tm8369* allele (Figure 2.B). Of note, these truncation alleles still carry the well-conserved PP2C domain (Figure 2.B). Therefore, we also generated a deletion null allele of *ppm-1.D* (*jf120)* (Figure 2.B). We validated this allele as a null by qRT-PCR (Figure S2.A). Both the truncation and null alleles displayed a small increase in embryonic lethality originating both from defective oogenesis and spermatogenesis (Figure S2.B and C). At very low frequency (2.6 ± 1.0 %, mean ± SD, n=1914), homozygous null *ppm-1.D* mutants sired progeny with abnormal body morphology indicating developmental defects (Figure S2.D).

Immunodetection of the tagged PPM-1.D (see Table S1 for functionality of the tagged lined) revealed a strong nuclear signal throughout the progenitor zone, which disappeared as soon as cells entered meiosis (Figure 2.C, top). The nuclear signal displayed a marked intensity increase at the nuclear periphery. In the proximal germline, PPM-1.D signal reappeared in diplonema as foci (Figure 2.C, middle) and later on a strong nuclear signal with enrichment at the nuclear periphery can be seen at diakinesis (Figure 2.C, bottom). The human ortholog PPM1D was reported as being expressed in response to p53 induction (Fiscella et al., 1997). CEP-1 (worm p53) is co-expressed in the germline progenitor zone (e.g., (Dello Stritto et al., 2021)), we therefore examined tagged PPM-1.D in the *cep-1* mutant (Figure S3.A). PPM-1.D expression was independent of *cep-1* in the germline. To test whether PPM-1.D is a substrate of the SCF^PROM-1^ ubiquitin ligase for targeted protein degradation, we examined the localization of PPM-1.D in the *prom-1(ok1140)* deletion background. In the *prom-1(ok1140)* mutant, PPM-1.D failed to disappear at meiotic entry and was detected at all stages of meiotic prophase (Figure 2.D), suggesting scheduled degradation of PPM-1.D by SCF^PROM-1^.

To test whether PPM-1.D is a direct PROM-1 substrate, we took advantage of the yeast *Saccharomyces cerevisiae* containing the conserved SCF complex subunits, but lacking a PROM-1 homolog. When PPM-1.D or PROM-1 are individually expressed in yeast, each protein was readily detected by Western blot. However, as soon as PROM-1 and PPM-1.D were co-expressed, PPM-1.D levels were significantly reduced (Figure 2.E, left). Addition of the proteasome inhibitor MG132 to cells co-expressing PROM-1 and PPM-1.D led to a 6 fold increase in PPM-1.D levels (Figure 2.E, right) reinforcing that the observed reduction of PPM-1.D is due to PROM-1 mediated degradation. This finding supports the idea that PPM-1.D is a target of the SCF^PROM-1^ complex.

PPM-1.D is a conserved protein, well known for its role in the response to DNA damage in mammals (Shaltiel et al., 2015). Here, we identify a novel activity, at the stage of meiotic entry, when meiotic progenitor cells differentiate. PPM-1.D has to be degraded by SCF ^PROM-1^ to mediated scheduled meiotic entry.

### CHK-2 and PPM-1.D are found together in protein complexes

As deleting *ppm-1.D* significantly rescues the meiotic *prom-1* mutant phenotypes and PPM-1.D is mostly expressed in the progenitor zone, we used endogenously tagged *ha::ppm-1.*D to determine the PPM-1.D interactome. Biochemical fractionation of germline cells revealed that PPM-1.D was enriched in the nuclear soluble and insoluble fractions (Figure 3.A). This is in agreement with our cytological analysis that PPM-1.D is detected in the nucleoplasm and is enriched at the nuclear rim (Figure 1.B).

**Figure 3.**
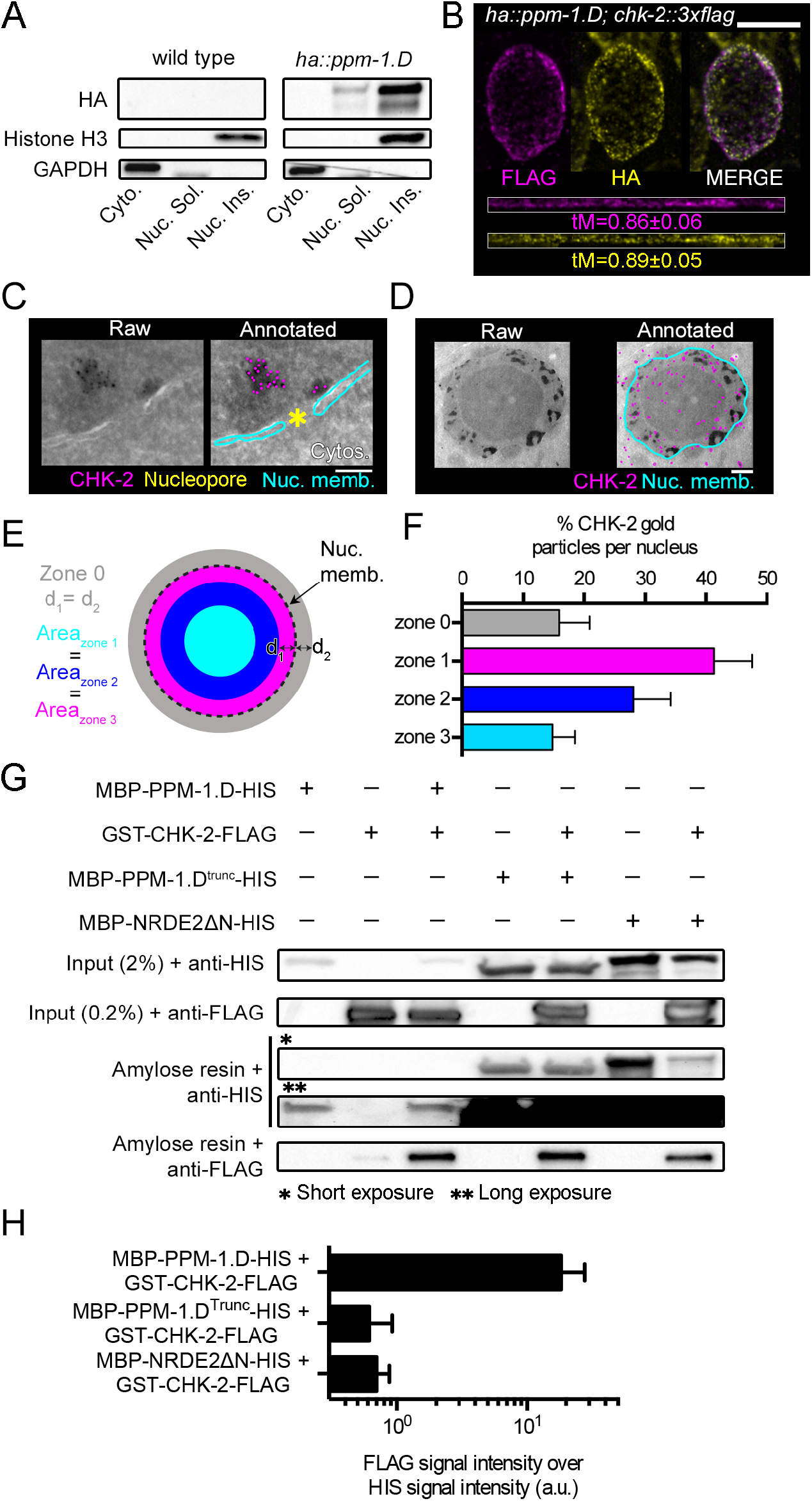
PPM-1.D and CHK-2 co-localize in the progenitor zone and interact physically. **A**. Western blot of cellular fractions (cytosolic, nuclear soluble and nuclear insoluble) with the specified antibodies for the indicated genotypes. **B**. STED visualized immuno-staining of CHK-2::3xFLAG (magenta) and PPM-1.D::HA (yellow) (top); straightened profiles of the signals (bottom). tM: automatic threshold Manders colocalization coefficient. Scale bar: 5 µm. **C**. Left, raw electron microscope image of one nucleopore with gold particles detecting CHK-2 and right with annotated nuclear membranes (cyan); CHK-2 (magenta). Scale bar: 10 nm. **D**. Left, raw electron microscope image of one mitotic nucleus with gold particles detecting CHK-2 and right with CHK-2 (magenta) and the nuclear membranes (cyan). Scale bar: 100 nm. **E**. Scheme used to divide the nucleus in 3 zones of equal area (zones 1, 2, and 3) and the outer vicinity of the nucleus (zone 0). **F**. Distribution of the CHK-2 gold particles in the different zones. **G**. Top, Western blot analysis after amylose purification for the indicated proteins expressed in *E. coli*. Bottom, quantification of the FLAG signal (CHK-2) normalized by the HIS signal for the mentioned co-lysed samples (n=2 Western blots).

Next, triplicated immuno-precipitation pull-down experiments of HA::PPM-1.D using the pooled nuclear fractions followed by mass spectrometry analysis revealed CHK-2 as a reproducible consistent interactor (Table S2). CHK-2 is a key meiotic regulator, involved in controlling numerous prophase I events in *C. elegans* (MacQueen and Villeneuve, 2001). To confirm the top-listed PPM-1.D-CHK-2 interaction, we endogenously tagged CHK-2 with a HA tag at the carboxy-terminus (see Table S1 for functionality) and performed triplicated immunoprecipitation experiments, followed by mass spectrometry analysis. Consistently, PPM-1.D was found in protein complexes containing CHK-2 kinase as top hit (Table S2).

### PPM-1.D and CHK-2 reside inside the nucleus

Since PPM-1.D and CHK-2 were reciprocally found as prime interactors in co-immunoprecipitations, we asked whether PPM-1.D and CHK-2 would also reside in the same sub-cellular compartments *in vivo*; (comprehensive CHK-2 localization in the germline has not been reported to date). We generated a strain expressing both HA::PPM-1.D and CHK-2::3xFLAG (for functionality of the CHK-2::3x FLAG, see Table S1) and examined their co-localization using STED microscopy. In the progenitor zone, PPM-1.D and CHK-2 showed striking co-localization in the nucleus, where both proteins were enriched at the nuclear periphery (Figure 3.B) and showed a high degree of staining overlap (automatic threshold Manders coefficient: CHK-2 = 0.86 ± 0.06, PPM- 1.D = 0.89 ± 0.05, average ± SD, 4 nuclei).

To assess whether the enrichment of CHK-2 at the nuclear rim was inside or outside the nuclear membrane, we employed electron microscopy with immunogold labeling. After validating the specificity of the antibody (Figure S4), we focused on the nucleopores. In the cryo-sections from progenitor zone nuclei, CHK-2 was in the close vicinity of the nucleopore in 38% of cases (13 out of 34 nucleopores) (Figure 3.C). At this resolution CHK-2 was found highly enriched in the nucleus both at the nuclear rim and inside the nucleus and a smaller fraction was detected in the cytoplasm (Figure 3.D). To quantify the signal, we divided each nucleus in three zones of equal area (zones 1, 2 and 3, Figure 3.E) and a fourth zone (zone 0) that is equidistant from the nuclear membrane as the zone 1 and represents the vicinity just outside of the nucleus. In each zone, we counted the number of gold particles detected in progenitor zone nuclei (Figure 3.F, n=20 nuclei). CHK-2 was mostly nuclear: 84.1 ± 9.6% of the gold particles were inside the nucleus and enriched in zone one (41.3 ± 6.2 %), just interior to the nuclear membrane. We conclude that in germline progenitor zone nuclei, CHK-2 is inside the nucleus and PPM-1D and CHK-2 strongly co-localize at the nuclear periphery.

### PPM-1.D directly interacts with CHK-2

As PPM1.D and CHK-2 were found associated in complexes and shared the same territories inside the nucleus we tested whether the *C. elegans* proteins interacted directly. To this aim we constructed MBP-PPM-1.D-10xHIS and GST-CHK-2-3xFLAG and expressed these proteins in *E. coli*. Both proteins were expressed and detectable in the cell lysates (Figure 3.G, input lanes). Next, we subjected the cell lysates to pull-down assays using amylose beads. Amylose beads purified MBP-PPM-1.D-10xHIS (Figure 3.G, first lane, amylose resin + anti-HIS, long exposure), and GST-CHK-2-3xFLAG displayed weak unspecific binding to the beads (Figure 3.G, second lane, amylose resin + anti-FLAG). When independent cultures of MBP-PPM-1.D-10xHIS and GST-CHK-2-3xFLAG were co-lysed and subjected to pull-downs, MBP-PPM-1.D-10xHIS co-purified GST-CHK-2-3xFLAG reproducibly (Figure 3.G, third lane, amylose resin + anti-FLAG), which suggests that PPM-1.D and CHK-2 can directly interact.

Next, we examined the binding of the truncated PPM-1.D protein lacking the last two exons (corresponding to the *(tm8369* or *jf76)* alleles, further referred to as truncated PPM-1.D) (Figure 2.B). Truncated PPM-1.D appeared more stable and more strongly expressed than the full-length protein in *E. coli* (Figure 3.G, fourth lane, input, anti-HIS), and was very efficiently purified using amylose beads (Figure 3.G, fourth lane, amylose resin + anti-HIS, short exposure). When CHK-2 was co-lysed with truncated PPM-1.D, we could only pull down low levels of CHK-2, when compared to normalized amounts of protein with full length PPM-1.D (Figure 3.G, fifth lane, amylose resin + anti-FLAG and Figure 3.H).

We also tested nonspecific sticking of CHK-2 protein to the MBP affinity tag. We expressed the unrelated protein, human NRDE2, (10xHIS-MBP-3C-NRDE2ΔN) with a similar molecular weight as PPM-1.D. After validating that we could efficiently purify 10xHIS-MBP-3C-NRDE2ΔN (Figure 3.G, sixth lane, short exposure, amylose resin + anti-HIS), we co-lysed GST-CHK-2-3xFLAG and MBP-3C-NRDE2ΔN-10xHIS expressing bacteria and performed MBP pull downs. We found that CHK-2 could be pulled down to similar levels with 10x-His-MBP-3C-NRDE2ΔN and MBP-3C-PPM-1.D-truncated-10xHIS (Figure 3.G, fifth and seventh lane respectively, amylose resin + anti-FLAG). As the truncated PPM-1.D and NRDE2 are both significantly higher expressed than the full-length PPM-1.D (Figure 3.G, first and third to seventh lane, compare short and long exposure, amylose resin + anti-HIS) and they both pull-downed similar amounts of CHK-2 (Figure 3.G, fifth and seventh lane, anti-FLAG), we conclude that CHK-2 was mostly binding to the MBP affinity tag, rather than to the PPM-1.D truncated protein.

Quantification of the amount of CHK-2 pulled down, normalized to input PPM- 1.D, revealed that CHK-2 binds full-length PPM-1.D 20 fold more efficiently than PPM- 1.D lacking the C-terminus, encoded by the two last exons (Figure 3.H, bottom, quantification derived from 2 biological replicates). We thus conclude that the PPM-1.D C-terminus is necessary for efficient interaction with CHK-2 or for protein folding to allow for efficient interaction.

### PPM-1.D restricts CHK-2 localization to the nuclear periphery

We first examined the pattern of CHK-2 and PPM-1.D localization in the progenitor zone and as germ cells enter meiosis. CHK-2 is expressed in the progenitor zone, overlapping with PPM-1.D (Figure 3.B). Sub-cellularly, CHK-2 shows strong co-staining with PPM-1.D at the nuclear rim, in the progenitor zone. In contrast, at and after meiotic entry the enrichment at the nuclear rim is lost, with CHK-2 being mostly nucleoplasmic in spots at the nuclear periphery (Figure 4.A), where it presumably co-localizes with putative substrates (e.g., the pairing center proteins, (Kim et al., 2015) or SUN-1 aggregates). We next examined CHK-2 localization in *ppm-1.D(jf120)* null and in the C-terminal truncation mutant *ppm-1.D* (*tm8369)*, which does not interact with CHK-2, both efficiently suppress the *prom-1(ok1140)* null phenotype (Figure 4.A). In both *ppm-1.D* mutant alleles, CHK-2 lost its nuclear rim enrichment in the progenitor zone and only nucleoplasmic signal was visible (Figure 4.A). These results are consistent with a model that PPM-1.D promotes the localization of CHK-2, in an inactive state, to the nuclear rim in progenitor zone cells, and that when PROM-1 degrades PPM- 1.D at meiotic entry, CHK-2 becomes nucleoplasmic and active. Furthermore, as *ppm- 1.D(tm8369)* results in loss of CHK-2 rim enrichment, we conclude that the C-terminal protein tail of PPM-1.D is necessary for CHK-2 enrichment at the nuclear rim in the progenitor zone.

**Figure 4.**
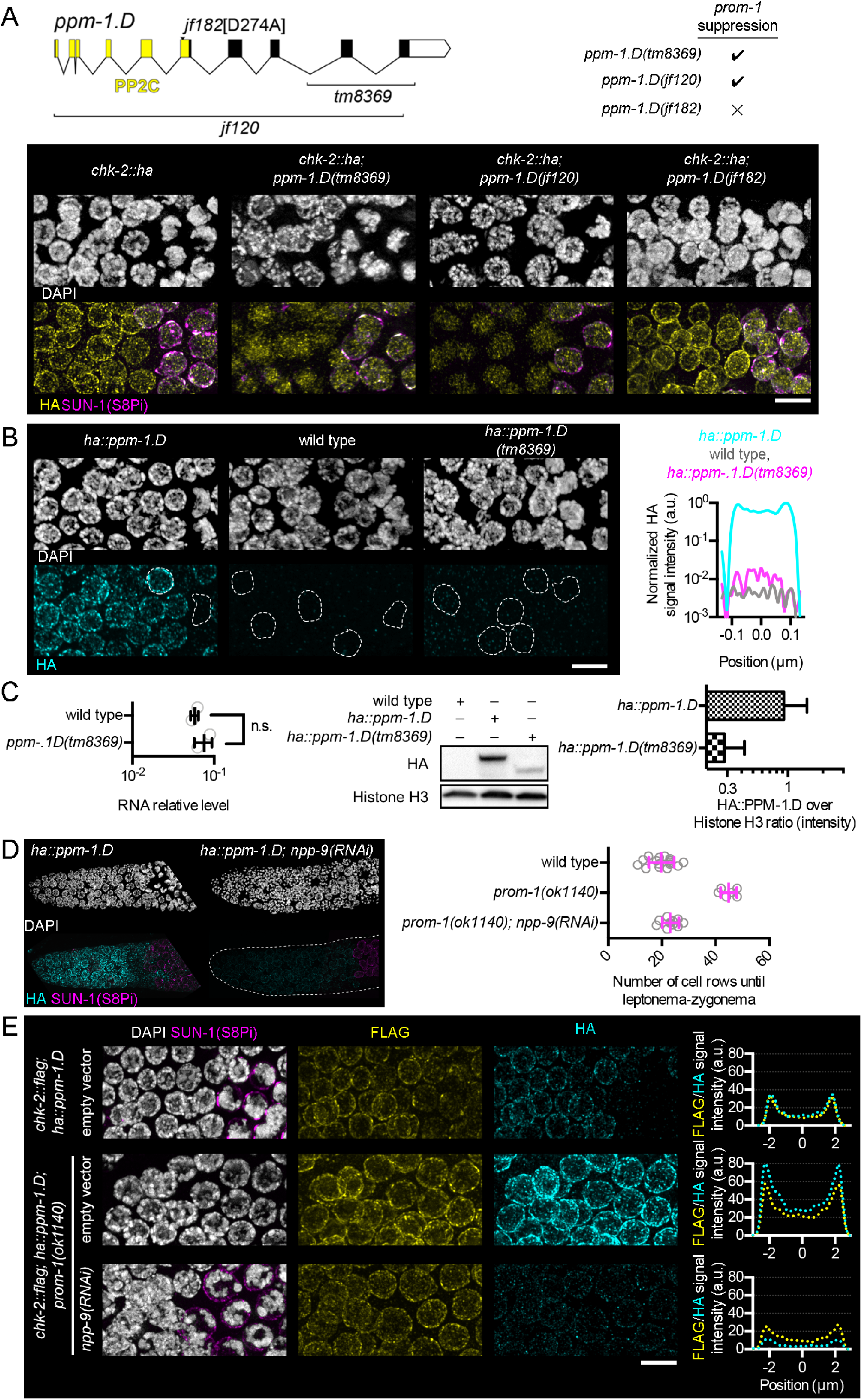
Regulation of CHK-2 localization and activity by PPM-1.D. **A**. Gene structure of *ppm-1.D* with domain, exon/intron structure and alleles (top, left), and genotypes suppressing *prom-1* phenotype. DAPI staining (white) and immuno-staining of HA (yellow) in the progenitor zone, for the indicated genotypes*. jf182* [D274A] allele is catalytic inactive PPM-1.D. Scale bar: 5 µm. **B**. DAPI staining and immuno-detection of HA (cyan) in the progenitor zone, for the indicated genotypes. Scale bar: 5 µm. Right, average line profile analysis of HA signal intensity centered on the nucleus for the mentioned genotypes (n=25 nuclei from the progenitor zone). **C**. Left, RNA quantification for *ppm-1.D*, for the indicated genotypes. Data for wild type is the same as in Figure S2.A. Center, western blot from whole worm extract for HA and the histone H3, for the indicated genotypes. Right, quantification of the ratio HA over histone H3 intensity, for the indicated genotypes. **D**. Left, DAPI staining and immuno-staining of PPM-1.D::HA (cyan) and SUN-1(S8Pi) (magenta) in the distal tip, for the indicated genotypes. Right, number of cell rows until entry into meiotic prophase, for the indicated genotypes. **E**. DAPI staining and immuno-detection of SUN-1(S8Pi) (magenta), FLAG (yellow) and HA (cyan) at the transition from progenitor zone to entry into leptonema-zygonema (centered at around 20 cell rows from the distal tip cell), for the indicated genotypes. Scale bar: 5 µm. Right, average line profile analysis of HA signal intensity centered on the nucleus, for the indicated genotypes (n=25 nuclei from the mitotic zone).

We then asked whether the catalytic activity of the PPM-1.D phosphatase was involved in the rim localization of both PPM-1.D and CHK-2. We mutated the aspartic acid (D) at position 274 to alanine to generate catalytic inactive PPM-1.D. D274 is highly conserved and part of the PP2C domain (Figure 4.A) and the exchange of aspartic acid to alanine was previously shown to abolish phosphatase catalytic activity (Takekawa et al., 2000). *ppm-1.D(jf182*[PPM-1.D(D274A)]) was validated as genetically inactive PPM-1.D (Figure S5), since addition of hydroxy urea resulted in equal levels of dead eggs as seen with the *ppm-1.D(jf120)* null allele. We next investigated the localization of CHK-2 in this mutant. Since CHK-2 nuclear rim staining was unaffected in *ppm-1.*D[D274A], we conclude that PPM-1.D catalytic activity was not required for the nuclear rim enrichment of CHK-2 in the progenitor zone. Remarkably, this catalytic inactive allele of *ppm-1.D* failed to rescue the *prom-1* phenotype (Figure 4.A).

We also sought to explore whether PPM-1.D localization to the nuclear periphery was dependent on CHK-2. Inactivation of *chk-2* with the allele *me64*, or deletion of the earlier identified paralogous gene, T08D2.7 (MacQueen and Villeneuve, 2001), corresponding to *chkr-2(ok431)*, or the double mutant, did not affect PPM-1.D nuclear rim staining (Figure S6). We therefore concluded that PPM-1.D nuclear periphery enrichment is *chk-2* and *chkr-2* independent. Summarizing, the sequestration of CHK-2 at the nuclear rim by PPM-1.D is independent of PPM-1.D phosphatase activity and CHK-2 activation does not require PPM-1.D phosphatase activity. Loss of PPM-1.D, via SCF^PROM^ mediated degradation, appears sufficient to liberate CHK-2 from the nuclear rim and allow the kinase to phosphorylate to initiate meiosis.

### PPM-1.D levels are regulating CHK-2

As the truncated allele of *ppm-1.D*, *tm8369*, retains the PP2C domain, we tagged the truncated protein to assess its expression. Truncated PPM-1.D displayed reduced nuclear staining without marked nuclear periphery enrichment when compared to the bright nuclear rim staining of wild-type PPM-1.D (Figure 4.B), reinforcing the idea that the C-terminal part of PPM-1.D is necessary for its nuclear periphery enrichment. Line profile analysis of the HA signal in *ha::ppm-1.D*-truncated across the nucleus showed that the detected signal is above the background level of the antibody measured on untagged worms (Figure 4.B, right). We then compared the mRNA levels of the full length and truncated *ppm-1.D* and this revealed that the mRNA of the truncated allele *ppm-1.D*(*tm8369)* is expressed at wild-type levels (Figure 4.C, left). We also quantified the levels of both wild type and the truncated HA::PPM-1.D by western blot (Figure 4.C, center), normalized to histone H3. The protein level of truncated PPM-1.D was three fold reduced when compared to the wild type (Figure 4.C, right). We therefore, conclude that the C-terminal part of PPM-1.D is necessary for protein stability. Moreover, we found that truncated PPM-1.D is still recognized by SCF^PROM-1^ for programmed/targeted degradation (Figure S7).

The loss of CHK-2 nuclear rim enrichment in the truncated allele (*tm8369*) could either be due to a reduction of PPM-1.D levels or due to the lack of the C-terminal part of PPM-1.D. To resolve the issue, we silenced the cytoplasmic nucleopore protein NPP-9 by RNAi to reduce the levels of PPM-1.D in the nucleus. This conditional knock-down of the nuclear pore gene *npp-9* led to a strong reduction of PPM-1.D staining in wild type, both in the nucleus and the nuclear rim. Moreover, silencing of npp-9 was able to reproducibly rescue the *prom-1* mutant phenotype (Figure 4.D). In *prom-1* mutants the leptonema-zygonema-like zone extends, on average 45 ± 3 (n = 6) cell rows from the distal tip of the germline, whereas in *prom-1; npp-9 RNAi* it extends 23 ± 3 (n = 10) cell rows, which is similar to wild type (20 ± 5 cell rows, n = 16).

We next examined PPM-1.D and CHK-2 localization in the *prom-1* mutant with and without *npp-9(RNAi)* to further comprehend this rescue. After *npp-9(RNAi)* PPM-1.D levels were reduced by three fold compared to wild-type levels (Figure 4.E, right) and at least seven times compared to *prom-1*. This three fold reduction was sufficient to promote scheduled meiotic entry as demonstrated by the timely phosphorylation of the CHK-2 substrate SUN-1 serine 8 (SUN-1(S8Pi), (Penkner et al., 2009). In addition, CHK-2 was localized both in the nuclear interior and nuclear rim associated. We conclude: 1) that the rim enrichment of CHK-2 is mediated though the C-terminal part of PPM-1.D and 2) that CHK-2 activity is responsive to the levels of PPM-1.D. Taken together, the C-terminal part of PPM-1.D is necessary for the localization of CHK-2 at the nuclear rim and the C-terminal truncation leads to instability of PPM-1.D. In addition the levels of PPM-1.D are regulating CHK-2 activity.

### Loss of PPM-1.D mediated CHK-2 inhibition leads to premature meiotic entry

PPM-1.D inhibits CHK-2. To promote meiotic entry, PPM-1.D is actively removed by SCF^PROM-1^ mediated proteolysis leading to activation of CHK-2, which is strongly correlated with relocation from the nuclear periphery to the nuclear interior. To test whether loss of PPM-1.D would lead to premature meiotic entry, we co-stained for CYE-1, a cyclin whose distal germline accumulation is restricted to the progenitor cell zone via SCF^PROM-1^ mediated proteolysis at meiotic entry (Biedermann et al., 2009; Fox et al., 2011), and SUN-1(S8Pi), a meiotic prophase marker for CHK-2 activity (Penkner et al., 2009) (Figure 5.A, top). These two markers show largely mutually exclusive accumulation, with nuclei expressing both markers were only rarely observed in wild type (Figure 5.A). Strikingly, in the *ppm-1.D* null allele, we found a consistent overlap of CYE-1 and SUN-1(S8Pi) accumulation in all germlines analyzed (Figure 5.A, bottom). We interpret this finding as SUN-1(S8Pi) appearing prior to downregulation of CYE-1, because of premature activation of CHK-2.

**Figure 5.**
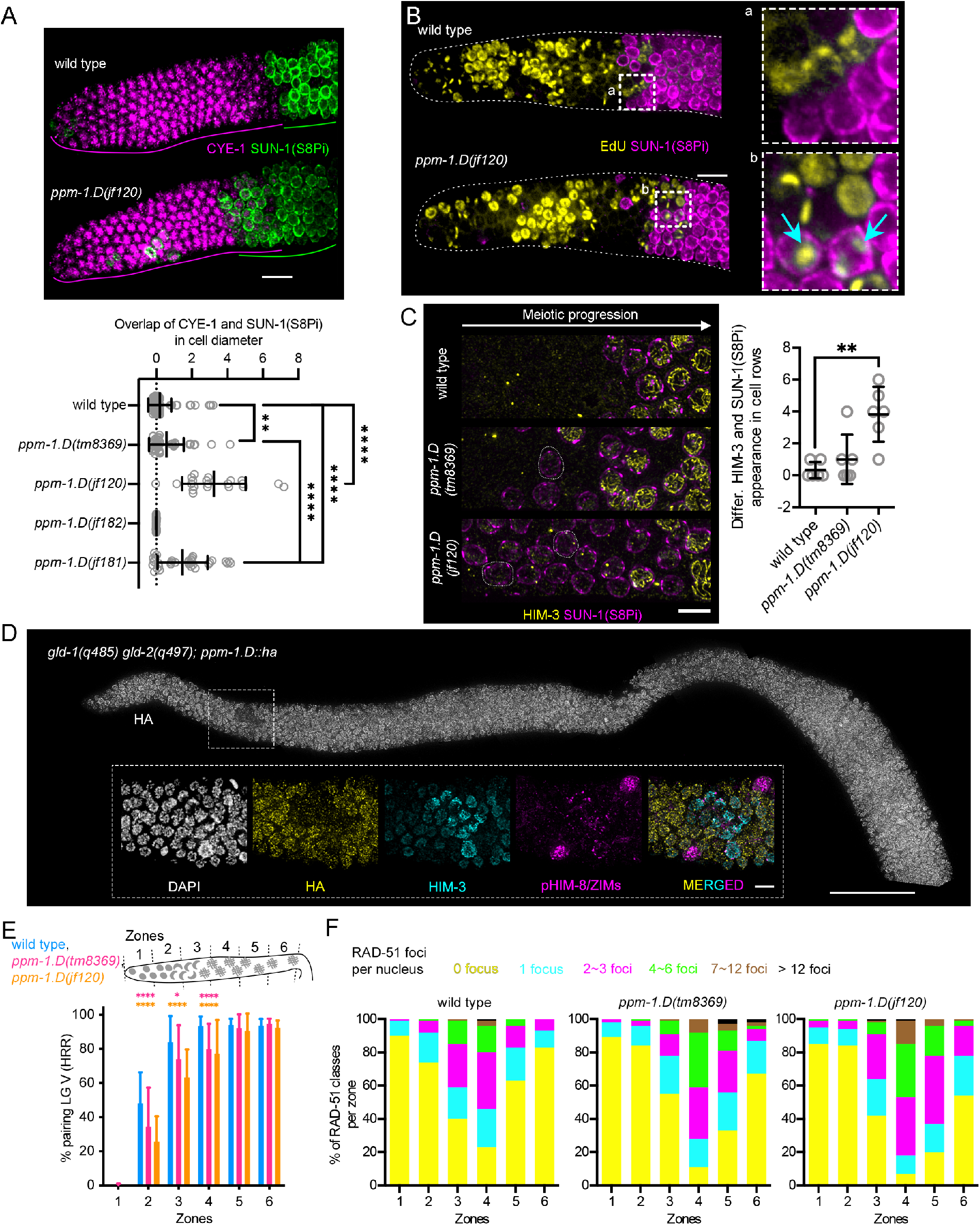
Premature meiotic entry in *ppm-1.D* mutants. **A.** Top, immuno-staining of CYE-1 (magenta) and SUN-1(S8Pi) (green) in the progenitor zone for the indicated genotypes. Scale bar: 10 µm. Bottom, distribution of the overlap between CYE-1 and SUN-1(S8Pi) staining in cell diameter, for the genotypes shown. **, P value <0.01, ****, P value <0.0001 for the Mann-Whitney test. **B**. Staining of EdU incorporation into replicating DNA (yellow) and SUN-1(S8Pi), for the indicated genotypes. Blue arrows in inset highlight nuclei with significant EdU incorporation, indicating ongoing meiotic S-phase, and staining for SUN-1(S8Pi), indicating CHK-2 activity and meiotic entry. Scale bar: 10 µm. **C**. Left, DAPI staining and immuno-staining of HIM-3 (yellow) and SUN-1(S8Pi) (magenta). Scale bar: 5 µm. Right, difference between the number of cell row at which HIM-3 and SUN-1 appears in the germline for the indicated genotypes. Cell rows were counted as positive when more than half of the cells were positive for the staining. **D**. Top, immuno-staining of HA in *gld-1(q485) gld-2(q497); ppm-1.D::ha* mutant worms. Scale bar: 50 µm. Insets show magnifications of nuclei stained for DAPI (white), HA (yellow), HIM-3 (cyan) and pHIM-8/ZIMs (magenta) in the zone highlighted in the top picture. Scale bar: 5 µm. **E**. Dissected gonads were divided into six zones of equal length. Percentage of nuclei with paired FISH signal; 5S probes on chromosome V in each zone, for the indicated genotypes. *, P value <0.05, ****, P value <0.0001 for the Fisher’s exact test. **F.** Percentage of nuclei with given number of RAD-51 foci in each zone, for the indicated genotypes. P values for the Fisher’s exact test are in Table S3.

We next examined staining in the *ppm-1.D* C-terminal truncation mutant, *tm8369*, also finding significant overlap of CYE-1 and pSUN-1 accumulation, although the extent of overlap was smaller than with the *ppm-1.D* null allele. Based on this difference, we hypothesize that both the catalytic activity and the C-terminal domain of PPM-1.D together contribute to CHK-2 inhibition/prevention of premature meiotic entry. To test this hypothesis, we mutated aspartic acid 274 (which leads to loss of catalytic activity) in the truncated *ppm-1.D* allele (intragenic double mutant *jf181*) and observed a significant increase in overlap between the two markers when compared to wild type or the C-terminal truncation allele (Figure 5.A, bottom). In contrast, removing only the catalytic activity of PPM-1.D did not lead to overlap between the markers. These results are in agreement with our previous observation that inactivation of the PPM-1.D catalytic domain alone was insufficient to rescue meiotic defects in *prom-1*. We propose that PPM-1.D exerts control over meiotic entry at two levels: 1) restraining CHK-2 localization to the nuclear periphery and 2) dephosphorylation of CHK-2 and perhaps other targets.

We next asked: what is the relationship between meiotic S-phase and meiotic entry in *ppm-1.D* null mutant? We monitored DNA synthesis by EdU incorporation into chromosomes (Kocsisova et al., 2018). In wild type, in a 30-min pulse labeling, EdU incorporation and SUN-1(S8Pi) staining are mutually exclusive. Significantly, in the *ppm-1.D(jf120)* mutant, some cells entered meiosis (SUN-1(S8Pi) positive cells) despite having replication still going on (EdU positive) (Figure 5.B). This phenotype was exclusively observed in the *ppm-1.D* null allele.

As our results suggest that in the absence of PPM-1.D CHK-2 is prematurely activated, we looked for possible direct consequences that could arise from premature CHK-2 induced meiotic entry. We reasoned that premature activation of CHK-2 might lead to uncoupling between meiotic chromosome axes formation, marked by HIM-3 loading (Zetka et al., 1999) and SUN-1(S8Pi). HIM-3 loading is independent of CHK-2, in contrast to the SUN-1 phospho-modification (Tang et al., 2010). Indeed, SUN-1(S8Pi) positive nuclei were observed in which HIM-3 had not assembled onto the chromosome axes, which is never the case in the wild type (Figure 5.C, left). The uncoupling between HIM-3 and SUN-1(S8Pi) was more prominent and significant in the *ppm-1.D* null allele (Figure 5.C, right).

To validate that lack of PPM-1.D is sufficient to activate CHK-2, we took advantage of the *gld-1(q485) gld-2(q497)* double mutant, which produces largely tumorous germlines with only very few cells entering meiosis, which eventually revert back to the progenitor fate (Mohammad et al., 2018). The few “meiotic cells” were devoid of PPM-1.D but showed expression of HIM-3 and CHK-2-mediated phosphorylation of the pairing center proteins (pHIM-8/ZIMs, (Kim et al., 2015)) (Figure 5.D). We conclude that the progenitor fate goes in hand with PPM-1.D presence and the loss of PPM-1.D correlates well with active CHK-2, whereas the CHK-2 independent loading of HIM-3 suggests that PPM-1.D regulates the activity of other targets.

We also examined the kinetics of chromosome alignment and pairing in the *ppm-1.D* mutants by FISH analysis using a probe for the 5S ribosomal RNA gene cluster. Pairing was delayed in both *ppm-1.D jf120 and tm8369* when compared to the wild type (Figure 5.E), however by pachynema the extent of pairing was indistinguishable from the wild type. Both *ppm-1.D* mutant alleles accumulated higher amounts of the marker of the meiotic recombination RAD-51 (Alpi et al., 2003; Colaiacovo et al., 2003), and a delayed clearance during the meiotic time course, which indicates an impediment of recombination. RAD-51 foci nonetheless disappeared, which suggests successful repair (Figure 5.F). In summary, we propose that meiotic entry in wild type occurs following the completion of meiotic S-phase and that premature meiotic entry, prior to completion of meiotic S-phase interferes with the kinetics of chromosome pairing and meiotic recombination. Further, we propose that both catalytic and non-catalytic activities of PPM-1.D together prevent premature meiotic entry.

### PPM-1.D is involved in the DNA damage response

DNA damage can stochastically appear during the mitotic cell cycle, and when it occurs a signaling mechanism induces repair to prevent aberrant cell divisions (Petsalaki and Zachos, 2020). In the *C. elegans* germline, DNA damage can occur in the progenitor zone caused by faulty mitotic replication or by random DNA insults, and after meiotic entry programmed DNA double strand breaks induced by the topoisomerase like enzyme SPO-11 (Dernburg et al., 1998). Persistent DNA damage will lead to an increased *cep-1*/p53-dependent apoptosis occurring at the end of pachynema (Gartner et al., 2008). We therefore set out to quantify apoptosis in the *ppm-1.D* mutants using SYTO12 as a reporter (Adamo et al., 2012). In comparison to the wild type, both *ppm-1.D* truncation (*tm8369*) and null (*jf120*) alleles displayed a significant increase in apoptotic corpses, indicating the presence of aberrant recombination intermediates (Figure 6.A). Deletion of *spo-11* in both *ppm-1.D* alleles failed to reduce the number of apoptotic corpses to wild type levels (Figure 6.A). Only the elimination of *cep-1/p53* in the *ppm-1.D* alleles led to the reduction of apoptosis to wild-type levels, supporting the view that *ppm-1.D* mutants accumulate both meiotic and *spo-11* independent DNA damage.

**Figure 6.**
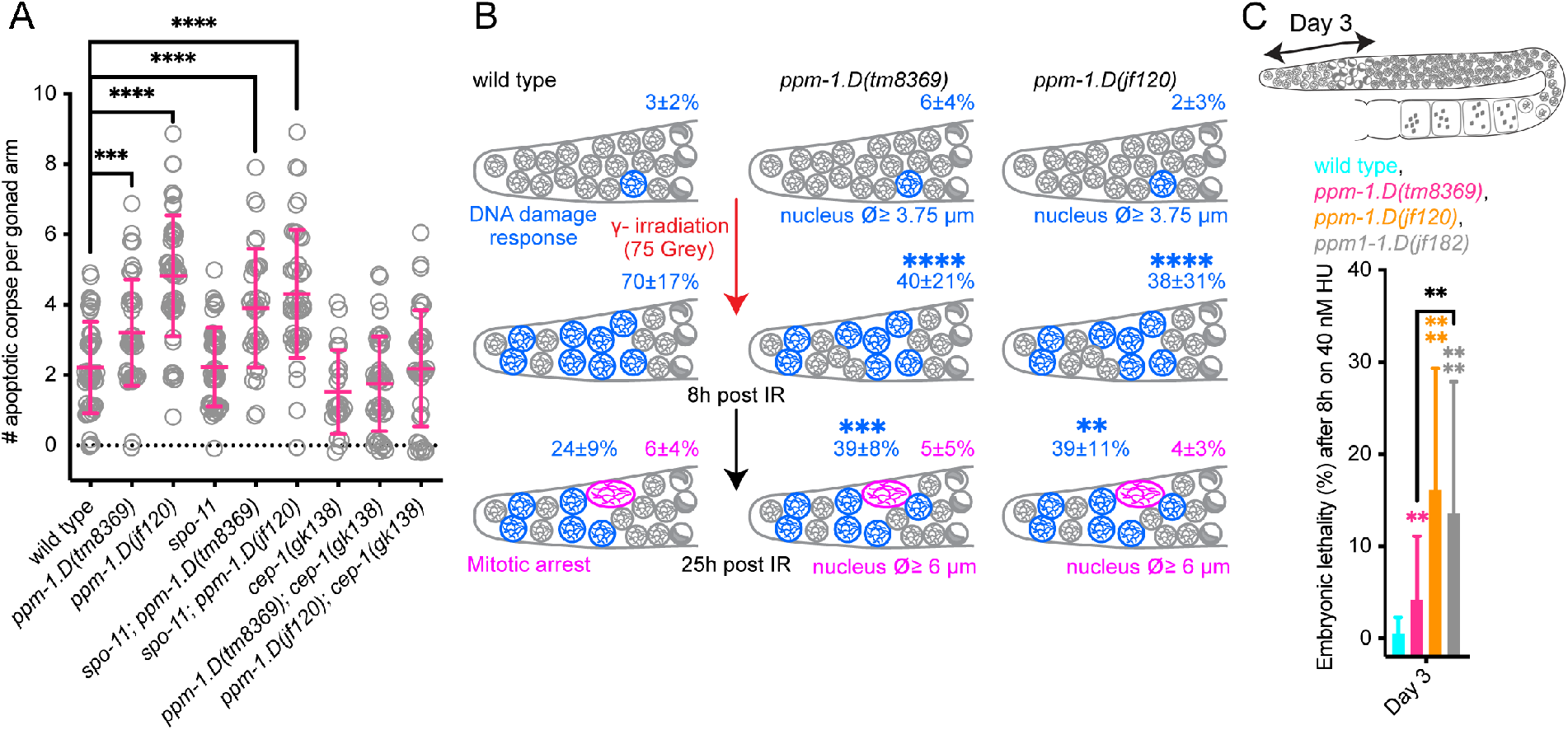
PPM-1.D functions in the DNA damage response. **A**. Quantification of apoptotic corpses (scatter and mean ± SD) for the indicated genotypes. ***, P value <0.001, ****, P value <0.0001 for the Mann-Whitney test. **B**. Percentage of nuclei with diameter above 3.75 µm before, 8 and 25 hours after γ-irradiation (75 grey), for the indicated genotypes. **, P value <0.01, ***, P value <0.001, ****, P value <0.0001 for the Fisher’s exact test. **C**. Embryonic lethality after 8 hours on 40 nM hydroxy urea (HU) three days after the stress, for the indicated genotypes. Data for wild type, *ppm-1.D(jf120)* and *ppm-1.D(jf182)* are the same as in Figure S5. The schematics of the *C. elegans* gonads (top) indicates the position of the nuclei in the germline at the time of exposure to irradiation. **, P value <0.01, ****, P value <0.0001 for the T test.

DNA damage could also arise in the mitotic germline compartment. We further challenged the DNA response in the progenitor zone by exposing worms to γ-irradiation (75Gy) (Figure 6.B) (Gartner et al., 2004b). In wild type, a response to DNA damage in this compartment leads to the enlargement of the nuclear diameter (Gartner et al., 2004a), which we measured 8 and 25 hours after γ-irradiation. Whereas in wild type after 8 hours, 70 ± 17% (average ± SD) of nuclei were responding to the challenge (nucleus diameter over 3.75 µm), both *ppm-1.D* alleles displayed a significantly slower activation (*ppm-1.D(tm8369)* 40 ± 21%, *ppm-1.D(jf120)* 38 ± 31%, average ± SD, Figure 6.B). We therefore concluded that PPM-1.D is promoting mitotic cell cycle DNA damage response. Further, we quantified mitotic M-phase arrest 25 h post γ-irradiation, which becomes evident as a nuclear diameter over 6 µm. Both *ppm-1.D* alleles lacked any significant increase in M-phase arrest compared to wild type and we conclude that PPM-.D was not promoting mitotic arrest. At this timepoint, *ppm-1.D* mutants still displayed a significantly increased number of enlarged nuclei (with a diameter over 3.75 µm): *ppm-1.D(tm8369)* 39 ± 8%, *ppm-1.D(jf120)* 39 ± 11%, average ± SD compared to the wild type 24 ± 9%, average ± SD (Figure 6.B). This might suggest that PPM-1.D is involved in DNA damage signaling either in induction and/or downregulation.

We also challenged progenitor zone cells by the depletion of nucleotides, which blocks DNA replication (Timson, 1975). After hydroxy urea (HU) exposure we measured embryonic lethality in wild type and *ppm-1.D* mutants. We focused on the lethality 3 days after the HU exposure, when exposed meiocytes were in the progenitor zone and early meiotic prophase. Both *ppm-1.D* alleles *tm8369* and *jf120*, displayed significantly increased lethality relative to the wild type (Figure 6.C). The *ppm-1.D* null allele led to a more severe embryonic lethality (day 3, Figure 6.C) than the C-terminal truncation allele, which is still expressed and contains a functional PP2C domain. As the catalytically inactive allele was as much affected as the null allele (Figure 6.C), we conclude that the lack of phosphatase activity was responsible for the increased lethality. Moreover, this implies that the phosphatase activity of the truncated allele is sufficient to partially rescue the stress induced by the replication block upon HU addition. Altogether our results show that, as in mammals, PPM-1.D is involved in the mitotic cell cycle response to DNA damage and to replication stress.

## Discussion

PPM-1D is a PP2C phosphatase and we isolated a recessive loss of function *ppm-1.D* allele in a screen aimed at suppressing the meiotic entry defects in the *prom-1* mutant. We found that like the mammalian protein (Shaltiel et al., 2015), PPM-1.D has the well-established canonical role in the DNA damage response. Importantly, we identified a novel function for PPM-1.D as a prominent factor involved in the transition from the progenitor cell fate to differentiation at meiotic entry. PPM-1.D is expressed in the germline progenitor zone cells and our data suggest that it is actively degraded by SCF^PROM-1^ at meiotic entry; indeed it seems a major target as evidenced by restoration of high levels of embryonic viability when suppressing *prom-1* defects. Our mass spectrometry data identified CHK-2 as the main interacting partner of PPM-1.D and we showed that the two proteins interact through the C-terminal domain of PPM-1.D. Moreover, we found that the C-terminal domain of PPM-1.D sequesters CHK-2 at the nuclear rim, promoting CHK-2 inactivation. Premature meiotic entry in *ppm-1.D* mutants leads to low levels of embryonic death, elevates rates of apoptosis, meiotic entry prior to completion of meiotic S-phase, the uncoupling of certain meiotic events (e.g. meiotic chromosome axes formation and chromosome end mobilization) and delayed chromosome pairing, which goes in hand with altered kinetics of meiotic recombination. *ppm-1.D* hermaphrodites sir progeny with developmental defects at a low rate, which could be explained by erroneous DNA repair taking place with a defective DNA damage response, but also if meiotic DSBs are induced prior to the completion of DNA replication.

### Control of meiotic entry in *C. elegans*

We propose the following model for meiotic entry in *C. elegans* (Figure 7). In the progenitor zone germ cells, PPM-1.D enters the nucleus, where it directly interacts with CHK-2 and sequesters CHK-2 to the nuclear periphery. This sequestration of CHK-2 at the nuclear rim depends on the C-terminal part of PPM-1.D protein and does not require its phosphatase activity. The rim co-localization represents the first layer of control of PPM-1.D over CHK-2. Interestingly, when we engineered a mutant lacking both the C-terminus and the catalytic activity (leaving the rest of the protein intact), we found a more pronounced premature meiotic entry than in the single mutants. We propose that both PPM-1.D mediated sequestration and phosphatase activity inhibit CHK-2 in the progenitor zone, although the sequestration maybe the predominant inhibitory mechanism. Meiotic entry is initiated via the programmed degradation of PPM-1.D. This scheduled degradation mediated by the SCF^PROM-1^ complex leads to the release of CHK-2 from the nuclear periphery which allows CHK-2 to successfully drive important processes during meiosis. CHK-2 antagonizing PPM-1.D activity appears to depend on its concentration. The amount of nuclear PPM-1.D may act like a toggle switch of CHK-2 activity, as suggested in *prom-1* mutant rescued by *npp-9* RNAi; here, only the nuclear amount of PPM-1.D was decreased, but CHK-2 remained nuclear periphery associated to a certain extent, however sufficient active CHK-2 was generated to rescue *prom-1*.

**Figure 7.**
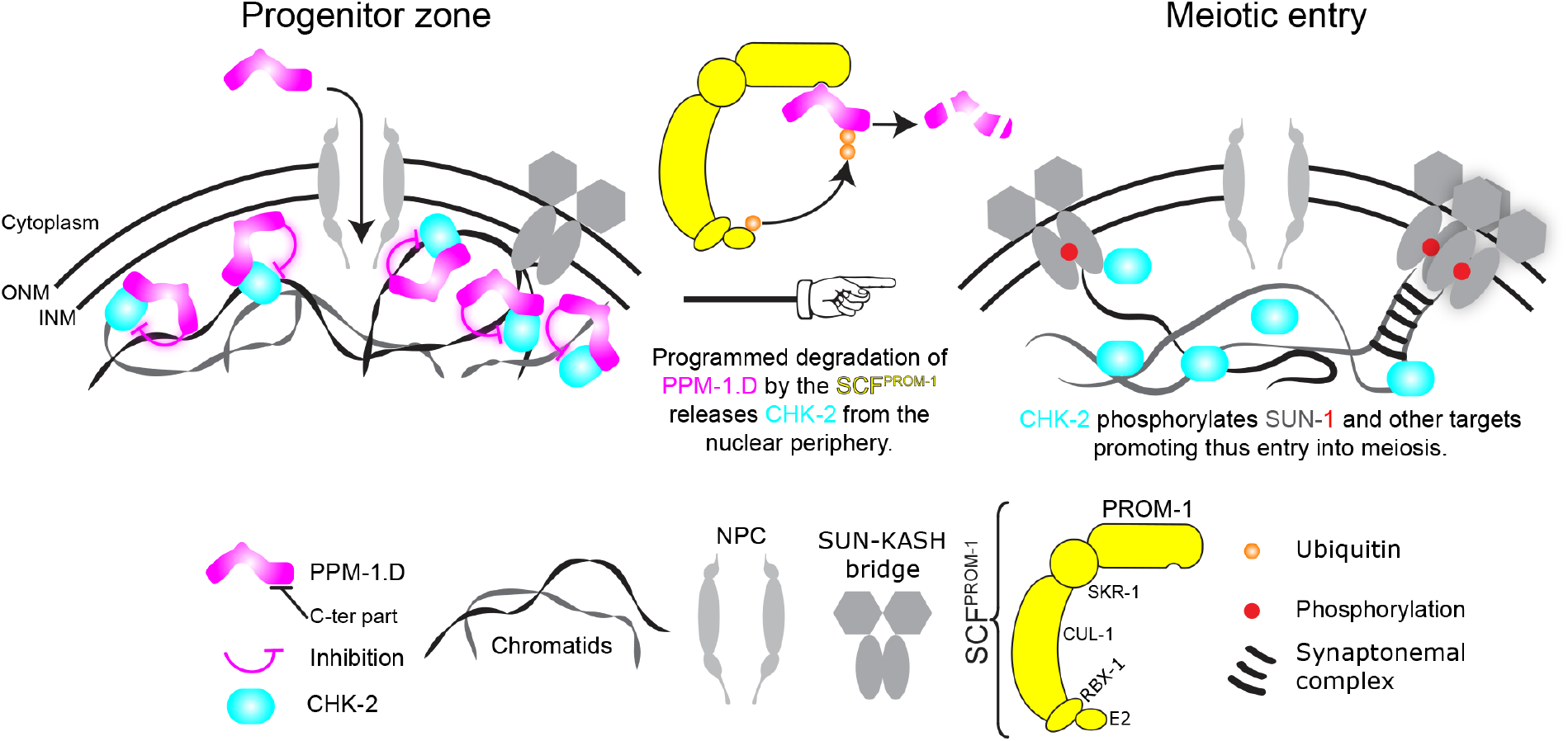
Model of control of meiotic entry by PPM-1.D. Entry of PPM-1.D into the nucleus is mediated by the nucleopores in the progenitor zone. Inside the nucleus PPM-1.D interacts directly with CHK-2 via its caboxy terminus and inhibits CHK-2 by sequestering it at the nuclear periphery and dephosphorylation. At meiotic entry, SCF^PROM-1^ degrades PPM-1.D. After the scheduled degradation of PPM-1.D, CHK-2 becomes released from the nuclear periphery, and gains access to its substrates and thus launches initial events of meiotic prophase.

### Dual function of PPM-1.D at meiotic entry

The PP2C phosphatase PPM-1.D first sequesters the meiotic key regulator CHK-2 (noncatalytic function) and second its phosphatase activity is involved in inactivating meiotic entry relevant targets (catalytic function), where CHK-2 could be one of several targets. The function of enzymes is not always restricted to their catalytic activity. For example, mammalian histone modifiers also exhibit noncatalytic roles involved in non-canonical processes like promoting cancer cell proliferation (Aubert et al., 2019), suggesting that enzymes having both noncatalytic and catalytic roles are potentially more common than expected. Similarly, there is growing evidence that phosphatases can lose their catalytic activity and gain non-catalytic activities through evolution (Reiterer et al., 2020). Such pseudo-phosphatases are involved in processes ranging from competition to substrate binding to spatial anchoring of binding partners. In the outlined scenario of meiotic entry PPM-1.D did not lose its phosphatase activity but exerts most of its control on CHK-2 via spatial sequestration of CHK-2 at the nuclear periphery, thus preventing premature meiotic entry.

### Regulation of CHK-2 by PPM-1.D and other potential targets

CHK-2 appears to be negatively regulated by PPM-1.D, however, there may be additional layers of regulation of CHK-2 in the progenitor zone. Indeed in *ppm-1.D* mutants, inappropriate activation of CHK-2 indicated by the premature appearance of SUN-1(S8Pi) is only confined to a couple of cell rows prior to meiotic entry and not to the entire progenitor zone. This could either mean that CHK-2 activation is regulated independently of PPM-1.D in the more distal region of the progenitor zone or that CHK-2 requires an activation step in addition to loss of inhibition by PPM-1.D. Moreover CHK- is potentially not the only target of the phosphatase PPM-1.D since the *prom-1* phenotype is more severe than the *chk-2* phenotype. *prom-1* mutants display defective cohesion and chromosome axes protein loading, which is not evident in *chk-2* mutants. PROM-1 has also been shown to function in the degradation of mitotic cell cycle proteins at meiotic entry (Mohammad et al., 2018). However, this function is not mediated by PPM-1.D (A. Mohammad, unpublished observations). Nevertheless, PPM-1.D is likely to function in the regulation of other meiotic proteins. We have observed similar defects in chromosome axes morphogenesis in *atr-1* (the worm ATL homolog) mutants (data not shown) thus PPM-1.D may also regulate the ATL-1 kinase at this important transition. Interestingly the uncoupling of chromosome axes loading and SUN-1 phospho-modification is less prominent in the *tm8369* truncation allele, which retains the catalytic activity of PPM-1.D. This could be a hint that the chromosome axes morphogenesis is predominantly under the control of the dephosphorylation activity of PPM-1.D.

### Conservation of the DNA damage response

In mammals, PPM1D/Wip1 is involved in the DNA damage response, the apoptotic response (Goloudina et al., 2016) and the protein is often overexpressed in cancer (Pechackova et al., 2017). In *C. elegans*, PPM-1.D is also involved in the response to DNA damage. Since PPM-1.D is also detected in the embryos (Figure 2.D - see embryo next to the progenitor zone tip), it would be very interesting to investigate its involvement in the regulation of the DNA damage response during developmental processes.

Upregulation of PPM1D/Wip1 expression in many human cancers makes the protein and attractive potential target for cancer therapy (Pechackova et al., 2017). It would be very interesting to determine whether the human homolog of PROM-1, FBXO47, specifically degrades PPM1D/Wip1. Renal carcinoma samples were identified with deletions in FBXO47 (Simon-Kayser et al., 2005), thus it would be highly interesting whether PPM-1.D/Wip1 qualifies as a target for FBXO47 as well and whether germline tumors are associated with mutations in FBXO47 in humans.

## Acknowledgments

We thank Stefan Schuechner, Marie Therese Kurzbauer, Luisa Cochella, Dea Slade, Anne Villeneuve, Nicola Silva, Monica Colaiacovo and Monique Zetka for sharing reagents, Egon Ogris and the Jantsch lab for fruitful discussions. We are indebted to Dieter Spittersberger and Angela Graf for their outstanding technical support and Nicolas Garcia-Seyda for strain construction. We are very thankful to Josef Gotzmann and Thomas Peterbauer for the state-of-the-art Microscopy Facility and valuable feedback. Some strains were provided by the CGC, which is funded by NIH Office of Research Infrastructure Programs (P40 OD010440). Electron microscopy was performed by the EM Facility of the Vienna Biocenter Core Facilities GmbH (VBCF), member of Vienna Biocenter (VBC), Austria. Mass spectrometry analysis was performed by the Mass spectrometry Facility of the Max Perutz Labs.

This project was funded by the Austrian Research Fund (FWF) project no. P 31275-B28 to V.J. T.S is supported by the National Institutes of Health grant R01 GM-100756. Y.K. is supported by the National Institutes of Health grant R35GM124895.

## Author Contributions

A.B., D.P., A.M., J.B. conducted and analyzed cell biology experiments;

A.B. performed the biochemistry and yeast analysis.

M.H. performed the mass spectrometry analysis.

R.L., S.F. performed the *E.coli* expression and purification.

A.B., D.P., J.B. constructed the worm strains;

A.B., D.P., A.M., S.F., T.S., Y.K. and V.J. conceived the project and analyzed data.

A.B., T.S and V.J. wrote the manuscript.

## METHODS DETAILS

### Nematode strains, strain construction, and culture conditions

All strains listed are derivatives of N2 Bristol and were cultivated under normal conditions (Brenner, 1974). Worms were γ-irradiated 24 hours post L4 stage with a dose of 75 Gy using a ^137^Cs source. CRISPR editing was done as described in (Paix et al., 2015) to the exception of prom-1::ha which was generated as described in (Norris et al., 2015). Guide and repair template as well as genotyping primers are listed in Supplemental Table S4.

### EMS screen

*prom-1(ok1140) unc-55(e402)* were grown on *E. coli* seeded plates for 5 days. On day 6, worms were collected in M9 buffer (0.3% KH_2_PO_4_, 0.6% Na_2_HPO_4_, 0.5% NaCl, and 1 mM MgSO_4_) and washed 3 times in M9 buffer to clean the worms from *E. coli*. Mutagenesis was carried out in 50 mM ethyl methane sulfonate (EMS). After mutagenesis, worms were allowed to recover until day 10 and then they were bleached to synchronize the population. L4 hermaphrodites were singled to small agarose plates seeded with *E. coli*. The viability of the mutagenized worms was assayed by looking for plates overcrowded in the second generation (F1+F2, Figure S1.A).

### Cytological preparation of gonads and immunostaining

Immunofluorescence was performed as previously described (Martinez-Perez and Villeneuve, 2005). L4 hermaphrodites were incubated at 20°C for 24 h. Gonads were then dissected from young adults into 1× PBS, fixed in 1% formaldehyde for 5 min at room temperature and frozen in liquid nitrogen. After post-fixation in ice-cold methanol, non-specific binding sites were blocked by incubation in PBS containing 1% BSA for at least one hour. Antibodies were diluted in 1x PBST (1x PBS, 0.1% Tween-20) and incubated overnight at 4°C (for primary antibodies) or 2 h at room temperature (for secondary antibodies). After washes in PBST, samples were mounted in Vectashield anti-fade (Vector Laboratories Inc., Burlingame, CA) containing 2 mg/ml 496-diamidino-2-phenylindole (DAPI).

For visualization of pHIM-8/ZIMs and HIM-3 (Figure 5.D) hermaphrodite germlines were dissected from 24 h post-L4 adults in egg buffer (25 mM Hepes, pH 7.4, 118 mM NaCl, 48 mM KCl, 2 mM EDTA, 5 mM EGTA, 0.1% Tween-20, and 15 mM NaN_3_) and fixed in 1% formaldehyde for 1 min before freezing in liquid nitrogen. Dissected germlines were further fixed in methanol at −20°C for 1 min and rehydrated with PBS with 0.1% Tween-20. Samples were then blocked with blocking reagent for 1 h and incubated with primary antibodies overnight at 4°C.

### RNA interference

RNAi was done as described in (Jantsch et al., 2004). Briefly, a single colony from the *npp-9* clone and the empty vector (Ahringer collection (Kamath et al., 2001)) were grown over-night at 37°C in 2xTY media supplemented with ampicillin (100 µg.ml^-1^). Next day cells were pelleted at 3,500 rpm for 15 min, resuspended in 2xTY and 150 µl of the suspension was used to seed NGM plates containing 1 M IPTG and 100 ngml^- 1^ampicillin. Bacterial growth was allowed at 37°C overnight. Pre-picked L4 were added to the plates and left at 20°C for 48h before analysis.

### RNA extraction and qPCR

Adult worms from 3 medium NGM plates were collected in M9 and allowed to sink in 1.5 ml Eppendorf tubes on ice. The supernatant was removed and 250 µl of Trizol was added and then the suspension was transferred to another 1.5 ml Eppendorf tube containing 150 µl of acid washed beads. Worms were broken open using a Fast Prep instrument (3 cycles: 15 s at 5,000g, 600 s rest). Mixture of broken worms was transferred into a new 1.5 ml Eppendorf tube. After addition of 50 µl of chloroform, samples were vortexed for 30 s and left at room temperature for 5 min. Next, samples were centrifuged at 12,000 rpm for 15 min at 4°C. The clear top layer was transferred into a fresh 1.5 ml Eppendorf tube and nucleic acids were precipitated by addition of 125 µl of isopropanol. Samples were spun down at 12,000 rpm for 10 min at 4°C. The pellet was washed with 500 µl of 70% ethanol and spun down at 14,000 rpm for 5 min at 4°C. The pellet was air dried and dissolved in 10 µl of RNAse-free water. After DNAse treatment using Promega kit following the provider instruction, cDNA synthesis was done using Superscript III with random hexamers as described in the kit. For the qPCR mastermix 100 ng of total RNA was used using the SensiFAST™ SYBR® No-ROX Kit and we used a Eppendorf Realplex 2 Mastercycler to read the plate. Ct measures were done in triplicate in the qPCR machine and these results were duplicated. *pmp-3* was used as reference (Zhang et al., 2012) and specific primers located in the 5’ and 3’ region of *ppm-1.D* were used to assess the RNA level. Results were analyzed using the delta-delta CT method (Schmittgen and Livak, 2008). Primers used are listed in supplemental Table S5.

### Microscopy and evaluation

3D stacks of images were taken using either a DeltaVision or a DeltaVision Ultra High Resolution microscope equipped with 100x/1.40 oil immersion objective lenses and a complementary softWORx software package. Images acquired with the DeltaVision where deconvolved using the softWORx deconvolution algorithm. Maximum intensity projections of deconvolved images were generated using ImageJ after adjustments of the maximums and background subtraction using a rolling ball radius of 50 pixels. Where specified, images of gonads consist of multiple stitched images. This is necessary due to the size limitation of the field of view at high magnifications. Stitching of images to build up entire gonads was performed manually in Adobe Photoshop. Levels of stitched images were adjusted to each other in Adobe Photoshop to correct for auto-adjustment settings of the microscope.

Super resolution images were acquired as single frame with an Abberior Instruments STEDYCON using alpha Plan-Apochromat 100x/1.46 Oil DIC with 2 avalanche photodiode detectors for dual-channel 2D STED (orange, dark red) with samples prepared as explained before except that samples were not mounted in DAPI but in Aberrior mounting media.

### Fluorescence in situ hybridization (FISH)

The FISH protocol is based on a published protocol (Silva et al., 2014). Dissected gonads were fixed in 4% paraformaldehyde in egg buffer for 2 min at room temperature and then stored in methanol at −20°C. Slides were then incubated in methanol at room temperature for 20 min, followed by 1 min washes in 50% methanol and 1× SCCT and dehydration by sequential immersion in 70%, 90% and 100% ethanol (3 min each). Hybridization mixture containing 10.5 μl FISH buffer (1 ml 20× SCCT, 5 ml formamide, 1 g dextran sulphate, 4 ml H2O) and 2.5 μl labeled probe was added to air-dried slides. The FISH probe for the 5S rDNA locus (chromosome V) was made by labeling 1 μg DNA with the DIG (Digoxigenin)-nick translation kit (Roche). After addition of EDTA, the probe was incubated at 65°C for 10 min. PCR-amplified 5S rDNA was used as probes the right end of chromosome V and was labeled by PCR with digoxigenin-11-dUTP. Slides were incubated at 37°C overnight in a humidified chamber and then washed twice (20 min) at 37°C in 50% formamide, 2Χ SCCT and 1Χ 10% Tween. After three washes in 2Χ SCCT at room temperature, samples were blocked for 1 h in 2Χ SCCT containing 1Χ BSA (w/v). Slides were then incubated in secondary anti-biotin antibody diluted in 2Χ SCCT (1:500) for 2 h at room temperature, followed by three washes in 2Χ SCCT, and then stained with 1 ng/ml DAPI and mounted in Vectashield.

### SYTO-12 Staining

Young adults (24 h post-L4 stage) were soaked in 33 μM SYTO-12 in PBS for 2–3 h at 20°C in the dark, transferred to unseeded NGM (nematode growth medium) plates for 30–60 min and then mounted. SYTO-12 positive cells were scored within the germline using an epifluorescence microscope equipped with a 40x or 63x oil immersion objective lens.

### Imaging and quantification of PROM-1 levels

Immunostaining was carried out as described (Mohammad et al., 2018). Briefly, synchronized 24-hr past L4, adult worms of the desired genotype are dissected in PBST (PBS with 0.1% Tween 20) with 0.2 mM levamisole to extrude the gonads. The gonads were fixed in 3% paraformaldehyde (PFA) solution for 10 min and then post-fixed in −20° chilled methanol for 10 min. After washing 3 x 10-min with PBST, they are blocked in 30% goat serum for 30 min at RT. The gonads are then incubated with the desired primary antibodies diluted in 30% goat serum at 4° overnight. Next day, after 3 x 10-min PBST washes, the gonads are further incubated with appropriate secondary antibodies, diluted in 30% goat serum, at 4° overnight. After three 10-min washes with PBST, the gonads were incubated with 0.1 g/ml DAPI in PBST for 30 min. After removal of excess liquid, the gonads were mixed with anti-fading agent (DABCO) and transferred to an agarose pad on a slide.

Quantification of PROM-1::HA was carried out similar to described (Chen et al., 2020), with some modifications. The dissected gonads were stained with primary antibodies against HA-tag, WAPL-1 and with DAPI. Hyperstack images are captured using a spinning disk confocal microscope (PerkinElmer-Cetus, Norwalk, CT). Exposure time for each channel were kept constant for an individual experiment. Two overlapping hyperstack images were captured to get a coverage of ∼50 cd from the distal end of the gonad. The images were further processed in Fiji and DAPI stained nuclei were used to mark the cell diameters (cds). Starting at the distal end, cd-wise plot profile (intensity) is extracted by using custom python script, for each gonad and are stored in text files. The intensity data was processed in R to visualize protein levels. Since PROM-1 quantification was carried out using antibodies against HA-tagged PROM-1, staining in N2, which lacks the HA-tagged PROM-1, was used to remove non-specific signals. WAPL-1 was used for the estimation of the progenitor zone length. All the scripts related to image processing and data analysis can be found at github (https://github.com/arizmohammad).

### Edu labelling

EdU labeling was carried out as described (Fox et al., 2011; Kocsisova et al., 2018; Mohammad et al., 2018). Briefly, synchronized 24-hr past L4 adult worms of the desired genotype were transferred to and fed on EdU-labeled plates for exactly 30 min before they were dissected and stained with the desired primary and secondary antibodies as described above. After overnight incubation with secondary antibodies, the gonads were given three 10-min washes with PBST, and then incubated with the EdU-detection reaction mix for 30 min at RT, using an EdU-labeling kit (Invitrogen). The gonads were given three 20-min washes with PBST to reduce background signal of EdU-labelling. The gonads were then incubated with DAPI and transferred to the slide as above.

### PPM-1.D and CHK-2 bacterial expression and pull-downs

The cDNAs encoding *C. elegans* CHK-2 and PPM-1.D were cloned into homemade vectors (derivatives of pBR322) harboring kanamycin resistance resulting in GST-CHK- 2-(3xFlag) and MBP-PPM1D-His10 fusion constructs. For protein production, the CHK- 2 and PPM1d constructs were transformed into *E. coli* BL21(DE3) derivatives and cells were grown at 37°C in terrific broth medium supplemented with kanamycin. When the *E. coli* cultures reached an optical density at 600 nm (OD_600_) of 2, the temperature was reduced to 18°C, and after 1 hour, protein production was induced by the addition of 0.2 mM IPTG for 12-16 h at 18°C over night. The next day, cells expressing CHK-2 or PPM- 1.D were either harvested individually or to test the interaction between CHK-2 and PPM-1.D, CHK-2 and PPM-1.D expressing cultures were mixed 1:1 before harvesting by centrifugation. Cell pellets were resuspended in 2 ml lysis buffer (50 mM Sodium phosphate, 25 mM TRIS/HCl, 250 mM NaCl, 20 mM Imidazole, 10% (v/v) glycerol, 0.05% (v/v) NP-40, 5 mM beta-mercaptoethanol pH 7.5) per g wet cell mass. Cells were lyzed by ultrasonic disintegration, and insoluble material was removed by centrifugation at 21,000xg for 10 min at 4°C. For MBP pull-downs 500 µL supernatant was applied to 35 µL amylose resin (New England Biolabs) and incubated for two hours at 4°C. Subsequently, the resin was washed three times with 500 µL lysis buffer. The proteins were eluted in 50 µL lysis buffer supplemented with 20 mM maltose.

### TCA protein precipitation

Overnight grown yeasts were refreshed in 5 ml synthetic media -Leu-Trp at an OD_600_ =0.05 and grown until their OD_600_ reached 0.8. For samples with MG132, MG132 (final concentration 10 µM) was added when cells reached an OD_600_ around 0.6 and let grow until they reached 0.8. As addition of MG132 reduces the division time of the yeast samples with MG132 were processed independently of others to avoid the introduction of artifacts by keeping other samples on ice.

1.25 ml of 100% ice cold TCA (20% final concentration) was added and cells were harvested (5 min, 4,500 rpm, 4°C). Cells were washed with 1 ml of ice-cold 10% TCA and transferred into 1.5 ml Eppendorf tubes. Next, cells were spun (10 min, 13,000 rpm, 4°C). 200 µl of ice-cold 10% TCA and 200 µl of acid washed glass beads were added to the pellet. Cells were broken using a FastPrep-24 5G instrument (MP Biomedicals) with 3 cycles (6.5 m/s, 45 sec, pause 5 min, 4°C). The supernatant was transferred to a fresh Eppendorf tube and beads were washed 3 times with 200 µl of ice-cold 10% TCA and the washes collected together with the supernatant. Eppendorf tubes were spun for 10 min at 5,000 rpm, 4°C. The pellet was resuspended in 200 µl of GSD buffer (40 nM Tris/HCl pH06.8, 8 M urea, 5% SDS, 0.1 nM EDTA, 2% (v/v) ß-mercaptoethanol, traces of bromophenol). After addition of 25 µl of unbuffered 1 M Tris base samples were boiled (10 min) and spun down (5 min, 1,000 rpm) before loading 5 to 30 µl on the SDS gel.

### Whole worm extract

Pre-selected L4 worms (200 per genotypes and per assay) were left at 20°C for 24 hours. Adults were collected into 30 µl TE buffer (10 mM Tris, 1 mM EDTA, pH8) into a 1.5 ml Eppendorf tube. After the addition of 1x Laemmli the Eppendorf tubes were submitted to three cycles of freeze thawing.

### Nuclei isolation and protein fractionation from large *C. elegans* cultures

Nuclei isolation and cellular fractionation were done as in (Silva et al, 2014). Briefly, large cultures of *C. elegans* were prepared by seeding twenty 100 mm NGM plates with 1 ml of OP50 bacteria (obtained from resuspending 2 liters of an overnight *E. coli* culture in a final volume of 40 ml). Between 5,000 to 6,000 *C. elegans* embryos were added to each 100 mm plate, and the plates were incubated at 20°C for three days. Young adult worms were collected and transferred to 50 ml tubes by washing the plates with M9, and tubes were left on a rack for 15 minutes to allow the worms to pellet by gravity, at which time most of the M9 was removed and fresh M9 solution was added. This washing step was repeated 3 times. The final wash was performed using NP buffer (10 mM HEPES-KOH pH 7.6, 1mM EGTA, 10 mM KCl, 1.5 mM MgCl2, 0.25 mM Sucrose, 1 mM PMSF and 1 mM DTT) containing protease inhibitors and worms were pelleted by centrifugation at 600 g for 2 minutes. 1 ml of this worm pellet was used to isolate nuclei. To isolate nuclei, worms were broken using a cooled metal Wheaton tissue grinder and the resulting worm solution was filtered first using a 100 μm mesh, followed by a second filtration with a 40 µm mesh. The filtered solution was then centrifuged at 300 g for two minutes at 4°C, and the supernatant from this step, which contains nuclei, was further centrifuged at 2,500 g for 10 minutes at 4°C. The resulting supernatant was used as cytosolic fraction, while germ line nuclei were contained in the pellet. In order to separate the nuclear soluble and the DNA-bound protein fractions from these nuclei, we used a Qproteome Nuclear Protein Kit from Qiagen according to manufacturer’s instructions.

### Western Blot

Samples were prepared as follows: 50 µg of the cellular fraction were mixed with 1x Laemmli whereas for the proteins extract from yeast the same amount of proteins (based on their OD_600_ at the time they were collected) was loaded to each well.

Samples were run in 1× SDS-Tris-glycine buffer on a pre-cast 4%-20% TGX gels (BioRad). Proteins were transferred onto PVDF membrane (activated in methanol for 20 seconds) for 1 hour at 4°C at 100V in 1× Tris-glycine buffer containing 20% methanol. Membranes were blocked for 1 hour in 1× TBS containing 0.1% Tween (TBST) and 5% milk; primary antibodies were added to the same buffer and incubated over night at 4°C. Membranes were then washed in 1× TBST and incubated with the secondary antibody in TBST containing 5% milk for 1 hour at room temperature. After washing, membranes were incubated with WesternBright ECL (Advansta) and developed with a ChemiDoc system (BioRad).

### Mass Spectrometry

Proteins were eluted from the beads by 3 x 20uL 100mM glycine, pH2. Supernatants were collected and the pH was adjusted to alkaline by addition of 1M TRIS pH 8. Disulfide bridge were reduced by DTT at a final concentration of 10mM for 30 min at room temperature. Free thiols were then alkylatyed with iodo acetamide (IAA) at a concentration of 20 mM for 30min at RT in the dark. Excess IAA was quenched with half of the amount of DTT used for reduction. Proteins were digested with 300ng trypsin over night at 37°C. Digests were acidified adding TFA to a final concentration of 1%. Peptides were desalted on StageTips (Rappsilber et al., 2007)and further purified according to the SP2 protocol by Waas (Waas et al., 2019).

Peptide samples were separated on an Ultimate 3000 RSLC nano-flow chromatography system (Thermo Scientific Dionex), using a pre-column for sample loading (PepMapAcclaim C18, 2 cm × 0.1 mm, 5 μm) and a C18 analytical column (PepMapAcclaim C18, 50 cm × 0.75 mm, 2 μm; both Thermo Scientific Dionex), applying a linear gradient from 2 to 35% solvent B (80% acetonitrile, 0.1% formic acid; solvent A 0.1% formic acid) at a flow rate of 230 nl/min over 120 minutes. Eluting peptides were analysed on a Q Exactive HF-X Orbitrap mass spectrometer (Thermo Scientific). For the data-dependent mode survey scans were acquired in a mass range of 375–1,500 m/z with lock mass on, at a resolution of 120.000 at 200 m/z. The AGC target value was set to 3E6 with a maximal injection time of 60 ms. The 8 most intense ions were selected with an isolation width of 1.6 and 0.2 m/z offset, and fragmented in the HCD cell with a normalized collision energy of 28%. Spectra were recorded at a target value of 1E5 with a maximal injection time of 150 ms and a resolution of 30000. Peptides with unassigned charge state, a charge of +1 or > +7 were excluded from fragmentation. The peptide match feature was set to preferred and exclude isotope feature wwas enabled. Selected precursors were dynamically excluded from repeated sampling for 30 s.

Raw data were processed using the MaxQuant software package 1.6.0.16 (http://www.maxquant.org/) (Cox and Mann, 2008) searching against the uniprot reference database of *C. elegans* and a costum made database of common contaminants. The search was performed with full tryptic specificity and a maximum of two missed cleavages. Carbamidomethylation of cysteine residues was set as fixed, oxidation of methionine, phosphorylation on serine, threonine and tyrosine, and N-terminal protein acetylation as variable modifications—all other parameters were set to default. The match between run feature and the search for 2nd peptides was enabled. Results were filtered at protein and peptide level for a false discovery rate of 1%. The protein groups table was imported into Perseus 1.6.2.1 (Tyanova et al., 2016), reverse hits and contaminants were filtered out as well as hits with less than 2 valid LFQ values in at least 1 experimental group. Missing LFQ values were imputed by values from a normal distribution. Data were statistically analyzed with LIMMA (Ritchie et al., 2015).

### Electron microscopy

24 hours post L4 stage *chk-2::ha* worms were immersed in 2% paraformaldehyde and 0.2% glutaraldehyde (both EM-grade, EMS, USA) in 0.1 M PHEM buffer (pH 7) for 2h at RT, then overnight at 4°C. The fixed gonads were embedded in 12% gelatin and cut into 1 mm^3^ blocks which were infiltrated with 2.3 M sucrose overnight at 4°C. These blocks were mounted onto Leica specimen carrier (Leica Microsystems, Austria) and frozen in liquid nitrogen. With a Leica UCT/FCS cryo-ultramicrotome (Leica Microsystems, Austria) the frozen blocks were cut into ultra-thin sections at a nominal thickness of 60nm at −120°C. A mixture of 2% methylcellulose (25 centipoises) and 2.3 M sucrose in a ratio of 1:1 was used as a pick-up solution. Sections were picked up onto 200 mesh Ni grids (Gilder Grids, UK) with a carbon coated formvar film (Agar Scientific, UK). Fixation, embedding and cryo-sectioning as described (Tokuyasu, 1973).

Prior to immunolabeling, grids were placed on plates with solidified 2% gelatin and warmed up to 37 °C for 20 min to remove the pick-up solution. After quenching of free aldehyde-groups with glycine (0.1% for 15 min), a blocking step with 1% BSA (fraction V) in 0.1 M Sörensen phosphate buffer (pH 7.4) was performed for 40 min. The grids were incubated in primary antibody, rabbit polyclonal to hemagglutinin, diluted 1:200 in 0.1 M Sörensen phosphate buffer over night at 4°C, followed by a 2h incubation in the secondary antibody, a goat-anti-rabbit antibody coupled with 6 nm gold, diluted 1:20 in 0.1 M Sörensen phosphate buffer, performed at RT. The sections were stained with 4% uranyl acetate (Merck, Germany) and 2% methylcellulose in a ratio of 1:9 (on ice). All labeling steps were done in a wet chamber. The sections were inspected using a FEI Morgagni 268D TEM (FEI, The Netherlands) operated at 80kV. Electron micrographs were acquired using an 11 megapixel Morada CCD camera from Olympus-SIS (Germany).

### Quantification of gold particles

Pictures were stitched in Photoshop to assemble the nucleus. The nuclear diameter was measured vertically, horizontally and the two diagonals using ImageJ. From the 4 measurements, we extracted the radius, *r*_1_, of the nucleus. To compute the radius of the two circles inscribed in the nucleus and dividing the nucleus into 3 areas of equal size we used the following formulas: 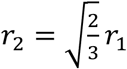 (radius of most outer inscribed circle) and 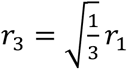 (radius of most outer inscribed circle). The nuclear membrane was traced in ImageJ with broken lines and using the line thickness the different zones were drawn. Gold particles were manually counted in Photoshop for each zone.

### Line profile analysis

Using ImageJ a line of 20 pixels width covering the diameter of a mitotic nucleus was created to measure the signal of HA antibody detection and added to the region of interest manager. At least 25 nuclei from the progenitor zone were processed this way. After collection of these line profiles, using R software the line profiles were resampled using the longest track as reference and then averaged. Averaged line profiles were plotted using GraphPad Prism6.

## QUANTIFICATION AND STATISTICAL ANALYSIS

Statistical analyses were performed in GraphPad Prism6. Datasets were tested for normal distribution; depending on outcome, populations were tested for significant differences using the two-tailed Fisher’s exact test or Mann–Whitney test or Chi-square test, as appropriate for each dataset.

## Strains and reagents

| REAGENT or RESOURCE | SOURCE | IDENTIFIER |
| --- | --- | --- |
| **Antibodies** |  |  |
| Rat anti HA (1:600) | Roche | Cat # 11867423001 |
| Rabbit polyclonal anti HA (1:1,000) | Sigma | Cat # H6908 |
| Rabbit anti WAPL-1 (1:2,000) | Novus | Cat # 49300002 |
| Guinea pig polyclonal anti HTP-3 (1:500) | (Goodyer et al., 2008) | N/A |
| Rabbit anti SYP-1 (1:500) | Gift from Nicola Silva, Masaryk University, Czech Republic | N/A |
| Rabbit anti REC-8 (1:100) | (Pasierbek et al., 2001) | N/A |
| Guinea pig anti-SUN-1 (1:500) | (Penkner et al., 2009) | N/A |
| Mouse anti FLAG (1:1,000) | SIGMA | Cat # F3165 |
| Rabbit anti-HIM-3 (1:750) | Novus | Cat # 53470002 |
| Guinea pig anti-SUN-1(S8Pi) (1:750) | (Penkner et al., 2009) | N/A |
| Mouse anti CYE-1 (1:10) | (Brodigan et al., 2003) | N/A |
| Mouse anti-HA (1:500) | Thermo Fisher Scientific | Cat # 26183 |
| Chicken anti-HIM-3 (1:500) | (Hurlock et al., 2020) | N/A |
| Rabbit anti pHIM-8/ZIMs (1 ug/ml) | (Kim et al., 2015) | N/A |
| Goat anti rabbit Alexa Fluor 568 (1:400) | Invitrogen | Cat # A-11036 |
| Goat anti guinea pig Alexa Fluor 488 (1:400) | Invitrogen | Cat # A-11073 |
| Goat anti mouse Alexa Fluor 594 (1:500) | Invitrogen | Cat # A-11032 |
| Goat-anti-rabbit 6 nm gold | Aurion | Cat # 800.011 |
| Donkey anti-Mouse Alexa Fluor 488 (1:200) | Invitrogen | Cat # A-21202 |
| Donkey anti-Rabbit Alexa Fluor 555 (1:200) | Invitrogen | Cat # A-31572 |
| Donkey Anti-Chicken Alexa Fluor 647 (1:200) | Jackson ImmunoResearch | Cat# 703-605-155 |
| Anti-mouse Abberior STAR 635P (1:200) | Abberior | Cat # ST635P-1001 |
| Anti-rabbit Abberior STAR 635P (1:200) | Abberior | Cat # ST635P-1007 |
| Mouse monoclonal anti HA (1:1,000) | Cell Signaling | Cat # 2367S |
| Rabbit polyclonal anti histone H3 (1:100,000) | Abcam | Cat # ab1791 |
| Mouse anti GAPDH (1:5,000) | Ambion | Cat # AM4300 |
| Mouse anti LexA (1:50,000) | Stefan Schuechner | N/A |
| Mouse anti FLAG (1:1,000) | SIGMA | Cat # F3165 |
| Goat anti rabbit HRP-conjugated (1:15,000) | Thermofisher | Cat # G21234 |
| Goat anti mouse HRP-conjugated (1:10,000) | Thermofisher | Cat # G21040 |
| **Chemicals, Peptides, and Recombinant Proteins** |  |  |
| Hydroxy Urea | Sigma-Aldrich | Cat # H8627-10G |
| Ethyl methane sulfonate (EMS) | Sigma-Aldrich | Cat # M0880-1G |
| (-)-Tetramisole hydrochloride | Sigma-Aldrich | Cat # L9756 |
| Blocking reagent | Roche | Cat # 11096176001 |
| **Critical Commercial Assays** |  |  |
| Qproteome Nuclear Protein Kit | Qiagen | Cat # 37582 |
| SYTO-12 | ThermoFischer | Cat # S7574 |
| Vectashield Mounting Medium | Vector Labs | Cat # H-1000 |
| EdU-labeling kit | Invitrogen | Cat # C10337 |
| amylose resin | New England Biolab | Cat # E8021S |
| Superscript III | Invitrogen | Cat # 18080051 |
| SensiFAST™ SYBR® No-ROX Kit | Bioline | Cat # BIO-98005 |
| **Experimental Models: Organisms/Strains** |  |  |
| *C. elegans*: N2 Bristol | CGC | https://cgc.umn.edu/strain/search |
| *C. elegans*: *prom-1(jf124[prom-1::ha])* | This study | UV145 |
| *C. elegans*: *ppm-1.D(jf76)III; prom-1(ok1140) unc-55(e402) I* | This study | UV157 |
| *C. elegans*: *ppm-1.D(tm8369)/qC1[dpy-19(e1259) glp-1(q339)] III* | This study | UV176 |
| *C. elegans*: *ppm-1.D(tm8369)III; chk-2(jf184[chk-2::ha]) V* | This study | UV177 |
| *C. elegans*: *ppm-1.D(jf120)/qC1[dpy-19(e1259) glp-1(q339)] III* | This study | UV178 |
| *C. elegans*: ppm-1.D(jf120) III; chk-2(jf184[chk-2::ha]) V | This study | UV179 |
| *C. elegans*: ppm-1.D(jf182[ppm-1.D(CD)]) III; chk-2(jf184[chk-2::ha]) V | This study | UV180 |
| *C. elegans*: ppm-1.D(jf181[ppm-1.D(tm8369+CD)]) III; chk-2(jf184[chk-2::ha]) V | This study | UV181 |
| *C. elegans*: chk-2(jf184[chk-2::ha]) V | This study | UV182 |
| *C. elegans*: ppm-1.D(jf183[ha::ppm-1.D]) III | This study | UV183 |
| *C. elegans*: chk-2(jf185[chk-2::3xFLAG]) V; ppm-1.D(jf183[ha::ppm-1.D]) III | This study | UV184 |
| *C. elegans*: prom-1(ok1140) unc-55(e402) I/ hT2[bli-4(e937) let-?(q782) qIs48](I;III) | This study | UV175 |
| *C. elegans*: ppm-1.D(jf183[ha::ppm-1.D]) III; prom-1(ok1140) unc-55(e402) /hT2[bli-4(e937) let-?(q782) qIs48](I;III) | This study | UV185 |
| *C. elegans*: ppm-1.D(jf183[ha::ppm-1.D]) III; prom-1(ok1140) unc-55(e402) /hT2[bli-4(e937) let-?(q782) qIs48](I;III); chk-2(jf185[chk-2::3xFLAG]) V | This study | UV238 |
| *C. elegans: spo-11(ok79)/nT1[unc-?(n754) let-? qIs50](IV;V).* | (Dernburg et al., 1998) | AV106 |
| *C. elegans: ppm-1.D(tm8369) III; spo-11(ok79)/nT1[unc-?(n754) let-? qIs50](IV;V).* | This study | UV186 |
| *C. elegans: ppm-1.D(jf120) III; spo-11(ok79)/nT1[unc-?(n754) let-? qIs50](IV;V).* | This study | UV187 |
| *C. elegans: cep-1(gk138) I* | (Hofmann et al., 2002) | TJ1 |
| *C. elegans: cep-1(gk138) I; ppm-1.D(tm8369)/qC1[dpy-19(e1259) glp-1(q339)] III* | This study | UV188 |
| *C. elegans: cep-1(gk138) I; ppm-1.D(jf120)/qC1[dpy-19(e1259) glp-1(q339)] III* | This study | UV189 |
| *C. elegans: ppm-1.D(jf183[ha::ppm-1.D]) III ; chk-2(me64) rol-9(sc148)/unc-51(e369) rol-9(sc148) V* | This study | UV190 |
| *C. elegans: ppm-1.D(jf183[ha::ppm-1.D]) III; chkr-2(ok431) X* | This study | UV237 |
| *C. elegans: ppm-1.D(jf183[ha::ppm-1.D]) III ; chk-2(me64) rol-9(sc148)/unc-51(e369) rol-9(sc148) V; chkr-2(ok431) X* | This study | UV191 |
| *C. elegans: fog-2(oz40)* | (Clifford et al., 2000) | BS553 |
| *C. elegans*: *gld-2(q497) gld-1(q485)/hT2 [bli-4(e937) let-?(q782) qIs48] (I;III) I; ppm-1.d::AID::HA (kim61) III; ieSi38 [sun-1p::TIR1::mRuby::sun-1 3'UTR + Cbr-unc-119(+)] IV* | This study | YKM393 |
| *E. coli* BL21(DE3) | Thermofisher | Cat # EC0114 |
| **Recombinant DNA** |  |  |
| Peft-3::cas9-SV40_NLS::tbb-2 3'UTR was a gift from John Calarco | (Friedland et al., 2013) | Addgene plasmid # 46168 |
| 3xHA::loxP::Pmyo-2_GFP::Prpl-28_neoR::loxP; gift from Monica Colaiacovo | (Norris et al., 2015) | N/A |
| Co-injection marker Pmyo-2::mCherry::unc-54utr; gift from Erik Jorgensen | (Frokjaer-Jensen et al., 2008) | pCFJ90<br>Addgene plasmid # 19327 |
| Co-injection marker pGH8 - pRAB-3::mCherry:: unc-54utr; gift from Erik Jorgensen | (Frokjaer-Jensen et al., 2008) | pGH8<br>Addgene plasmid # 19359 |
| Peft-3::Cre | (Dickinson et al., 2013) | pDD104<br>Addgene plasmid # 47551 |
| Homemade derivative of pBR322 (kanamycin resistance) GST-3C-CHK-2-(3xFlag) (nematode CHK-2) | This study | N/A |
| Homemade derivative of pBR322 (kanamycin resistance) MBP-3C-PPM1D-His10 (nematode PPM-1.D) | This study | N/A |
| Homemade derivative of pBR322 (kanamycin resistance) MBP-3C-PPM1D (truncated)-His10 (nematode PPM-1.D) | This study | N/A |
| Homemade derivative of pBR322 (kanamycin resistance) 10xHIS-MBP-3C-NRDE2ΔN (human NRDE2) | This study | N/A |
| **Oligonucleotides** |  |  |
| Guide, repair template and genotyping primers used in this study in 5' to 3' orientation as DNA sequences. | This study | Table S4 |
| Primer pair used for qPCR. | This study | Table S5 |
| **Software and Algorithms** |  |  |
| ImageJ | Schneider et al., (2012) | https://imagej.nih.gov/ij/ |
| Adobe Photoshop CC2018 | Adobe | N/A |
| Adobe Illustrator CC2018 | Adobe | N/A |
| GraphPad Prism 6 | GraphPad | N/A |
| RStudio | RStudio | https://www.rstudio.com |
| R | The R foundation | https://www.r-project.org/ |

## Supplemental information

### Supplemental figures and legends

**Figure supplemental 1.**
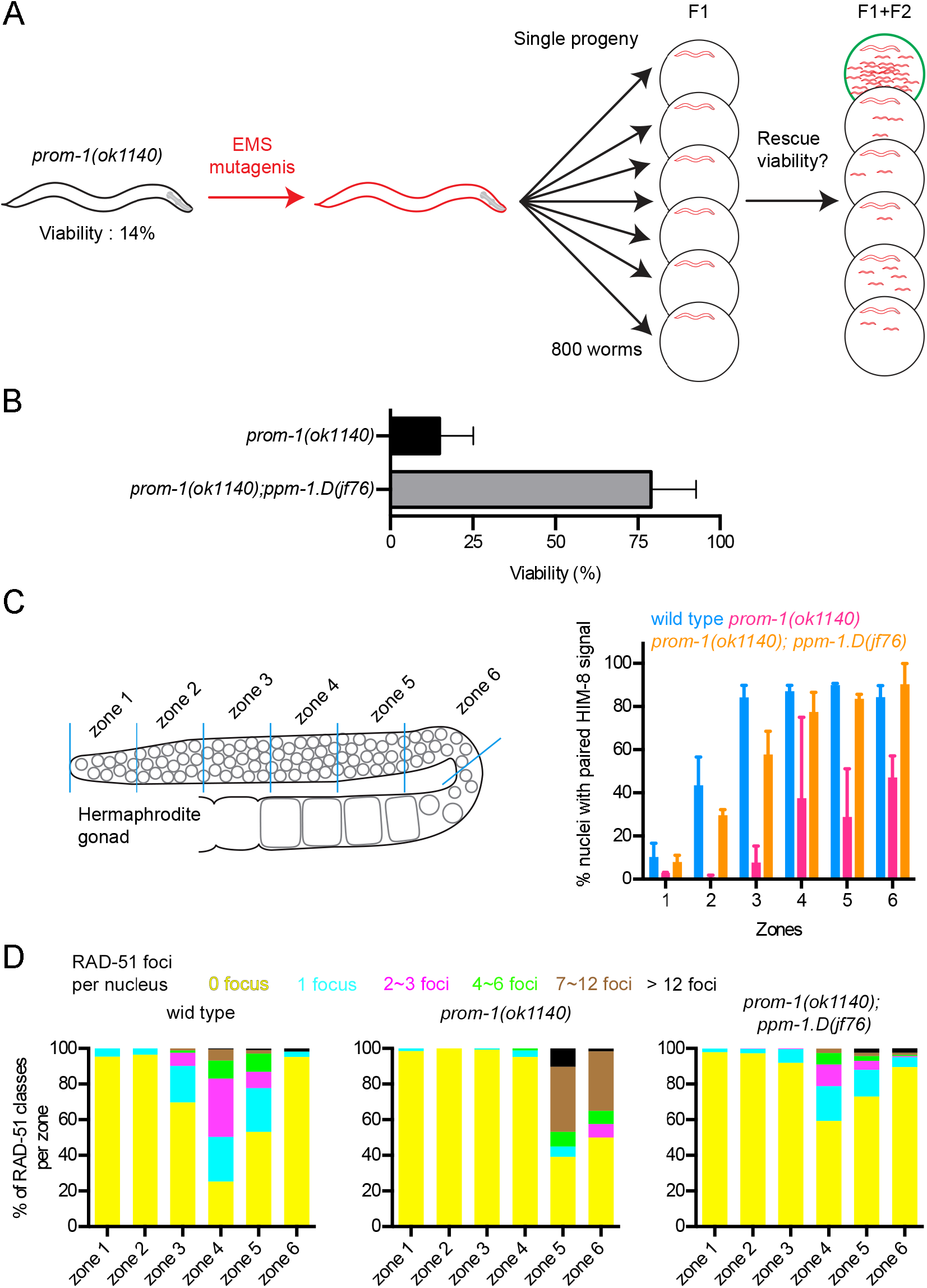
Identification of *prom-1* suppressor and characterization of the double mutant *prom-1(ok1140); ppm-1.D(jf76)*. A. schematics of the suppressor screen. F1 heterozygotes were singled after mutagenesis and suppressor candidate plates scores based on the viability/population density on the plates. B. Viability of *prom-1(ok1140)* and the suppressor line *prom-1(ok1140); ppm-1.D(jf76)*. C. Left: *C. elegans* hermaphrodite gonad divided into 6 zones of equal lengths. Right: percentage of X chromosome pairing (scored with HIM-8) in the different zones for the mentioned genotypes. D. Quantification of RAD-51 foci counted in the different zones for the mentioned genotypes.

**Figure supplemental 2.**
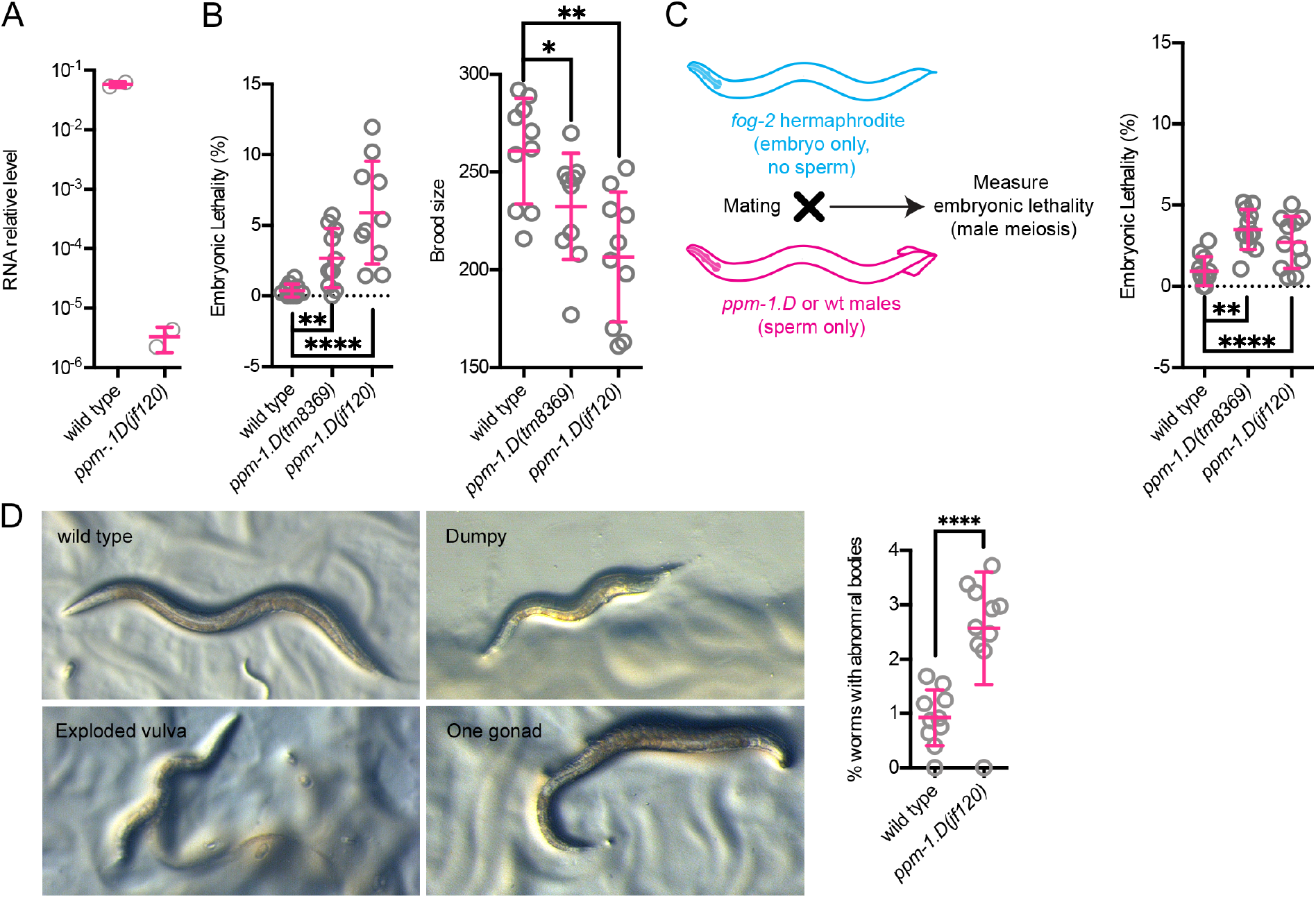
*ppm-1.d* mutants display low levels of unhatched embryos originating from both defects in oogenesis and spermatogenesis. **A**. Relative levels of *ppm-1.D* RNA in the mentioned genotypes. **B**. Left. Embryonic lethality in percentage for the mentioned genotypes. Right. Brood size counts for the mentioned genotypes. **C**. Left, *ppm-1.D* mutant males were mated to *fog-2* mutants to test male meiosis. Right, Percentage of non-hatching eggs for the mentioned genotypes. *, P value <0.05, **, P value < 0.01, ****, P value < 0.0001 for the Man-Whitney test. **D**. Left, representative pictures of abnormal body morphologies observed in *ppm-1.D(jf120)*. Right, quantification of abnormal *in* wild type and *ppm-1.D(jf120)* worms. 2000 synchronized worms were screened for abnormal body morphologies for each genotype. ****, P value < 0.0001 for the Chi-square test.

**Figure S3.**
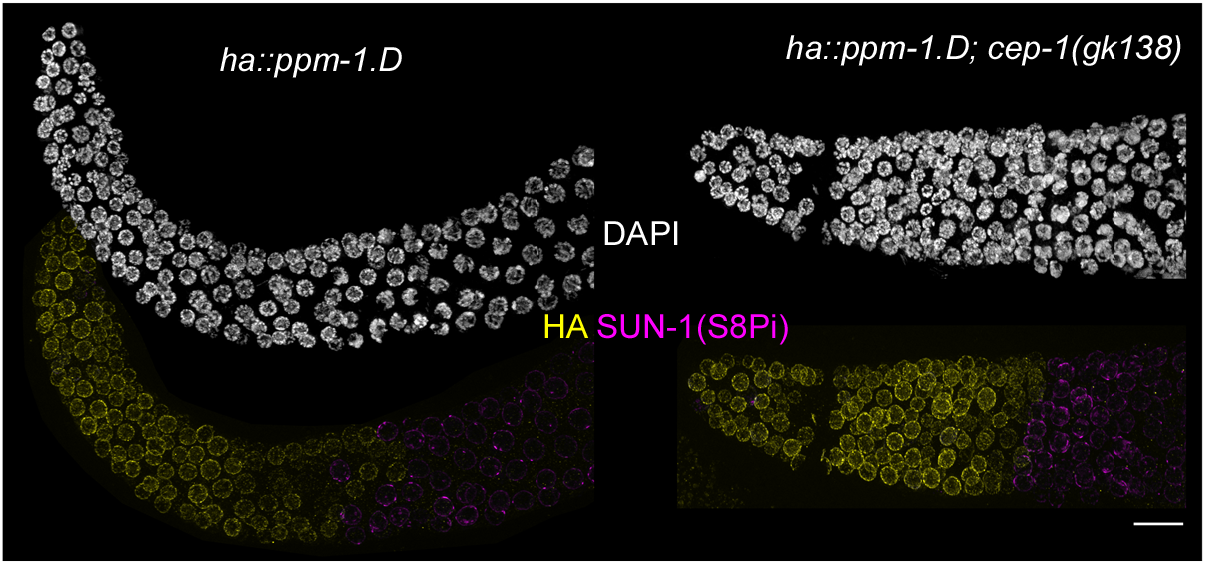
PPM-1.D expression in the progenitor zone of is not controlled by *cep-1.* DAPI staining and immuno-staining for HA::PPM-1.D (yellow) and SUN-1(S8Pi) (magenta) for the given genotypes. Scale bar: 10 µm

**Figure Supplemental 4.**
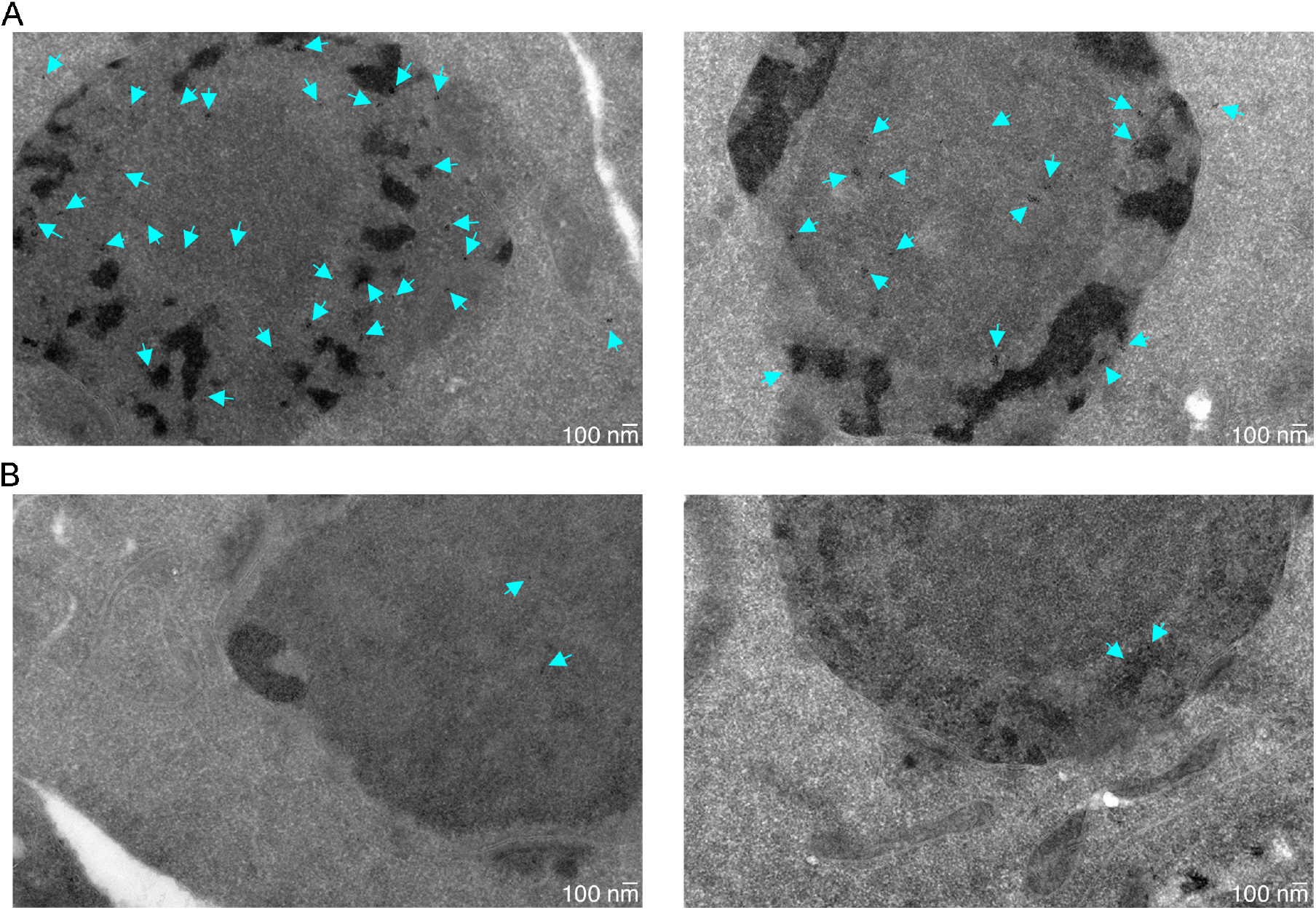
Specificity of the antibody used in electron microscopy. **A.** Representative pictures of mitotic nuclei at 14,000x resolution with cyan arrows highlighting the gold particles linked to the secondary antibody recognizing the primary antibody. **B**. Representative pictures of mitotic nuclei at 14,000x resolution with cyan arrows highlighting the gold particles linked to the secondary antibody without primary antibody.

**Figure Supplemental 5.**
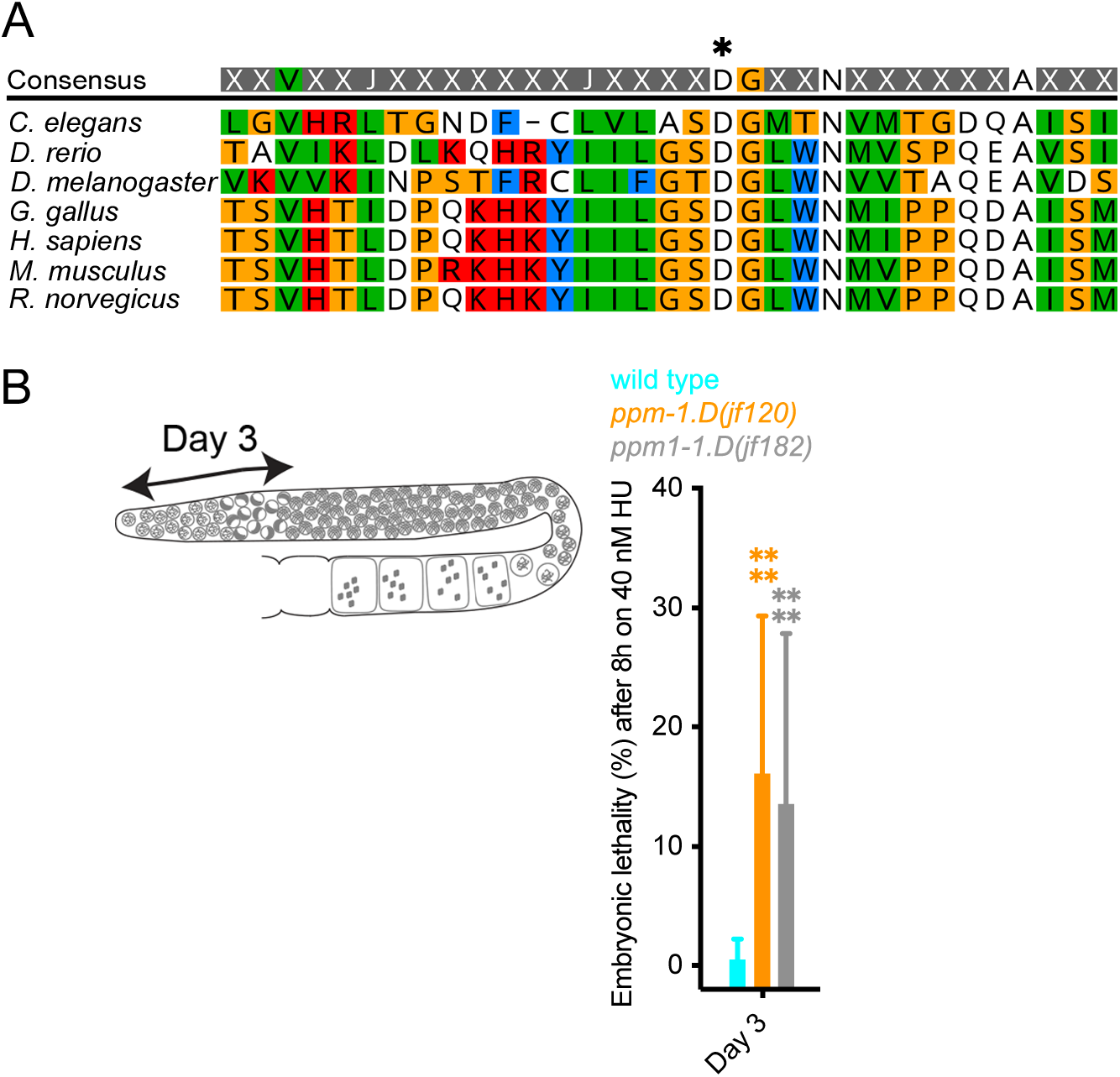
Validation of catalytic inactive PPM-1.D. **A**. Alignment of PPM-1.D protein sequences (amino acids 498 to 530) for the mentioned organisms highlighting the conservation of the PP2C domain. Asterisk marks the conserved aspartic acid required phosphatase activity (Takekawa et al., 2000). **B.** Scheme of *C. elegans* germline indicating the position of the nuclei in the gonad at the time of the irradiation and the day at which their embryonic viability can be measured. **C**. Embryonic lethality after 8 hours on 40 nM hydroxy urea 3 days after the stress for the mentioned genotypes. *jf120* allele is a null allele of *ppm-1.D* and *jf182* encodes catalytic inactive PPM-1.D. ****, P value <0.0001 for the Mann-Whitney test.

**Figure Supplemental 6.**
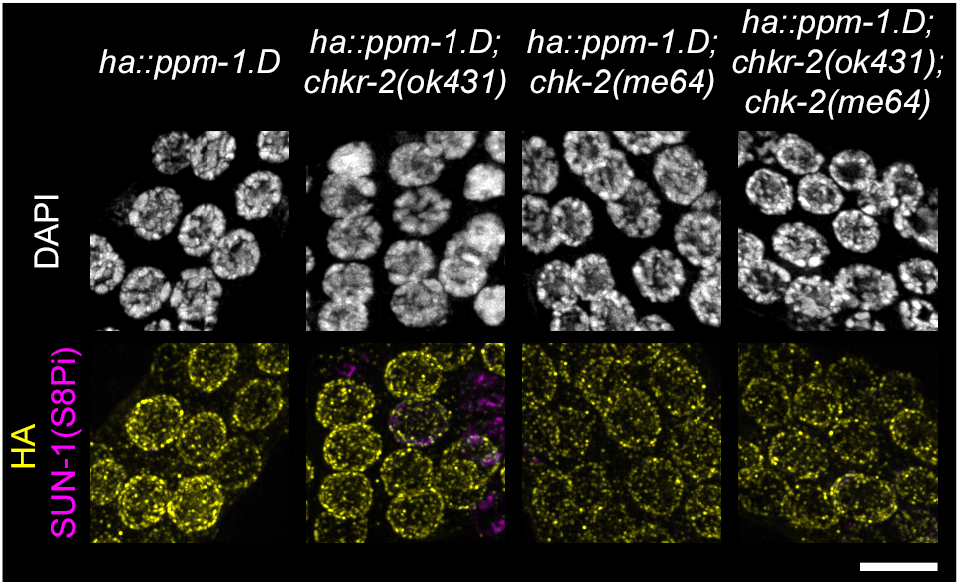
HA::PPM-1.D localization at the nuclear periphery is independent of *chk-2* and its paralog *chkr-2*. DAPI staining and immuno-staining of HA (yellow) and SUN-1(S8Pi) (magenta) for the given genotypes. Scale bar: 5 µm

**Figure supplemental 7.**
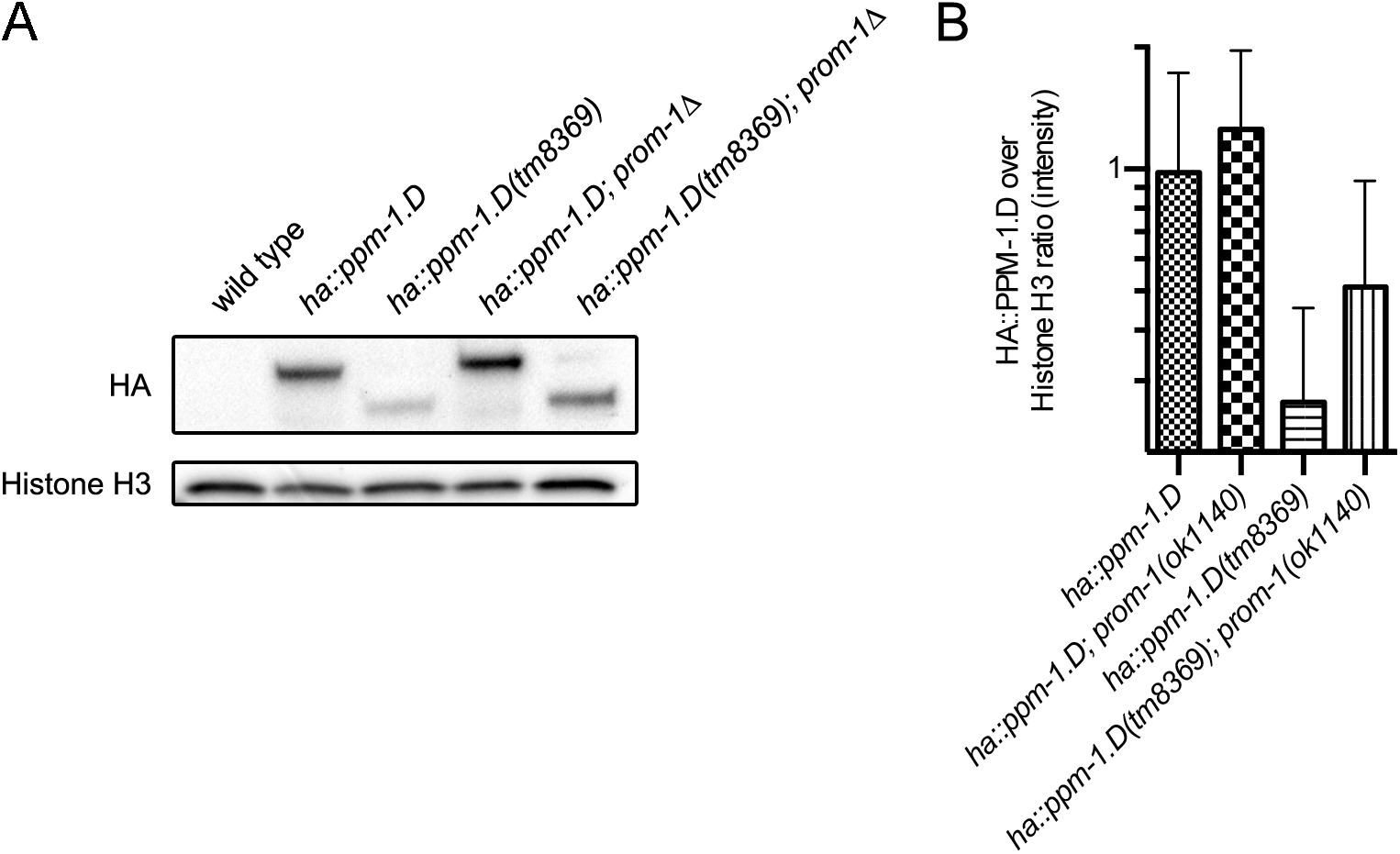
PPM-1.D^truncation^ is regulated by the SCF^PROM-1^ complex. **A**. Western blot from whole worm extracts for HA::PPM-1.D and the histone H3. **B**. Quantification of the ratio HA::PPM-1.D over histone H3 for the mentioned genotypes. Data for both wild type, *ha::ppm-1.D* and *ha::ppm-1.D(tm8369)* are the same as in figure 4C in **A** and **B**.

### Supplemental tables and legends

**Table S1.** Viability for the mentioned *C. elegans* strains. Progeny of 10 worms were scored.

| Strain genotype | Viability (% average ± SD) |
| --- | --- |
| **Wild type** | 99.74 ± 0.24 |
| *prom-1::ha* | 99.23 ± 0.60 |
| *ha::ppm-1.D* | 99.66 ± 0.41 |
| *ppm-1.D::ha* | 96.7 ± 1.70 |
| *chk-2::ha* | 97.97 ± 2.28 |
| *ha::ppm-1.D; chk-2::FLAG* | 99.84 ± 0.25 |

**Table S2.**
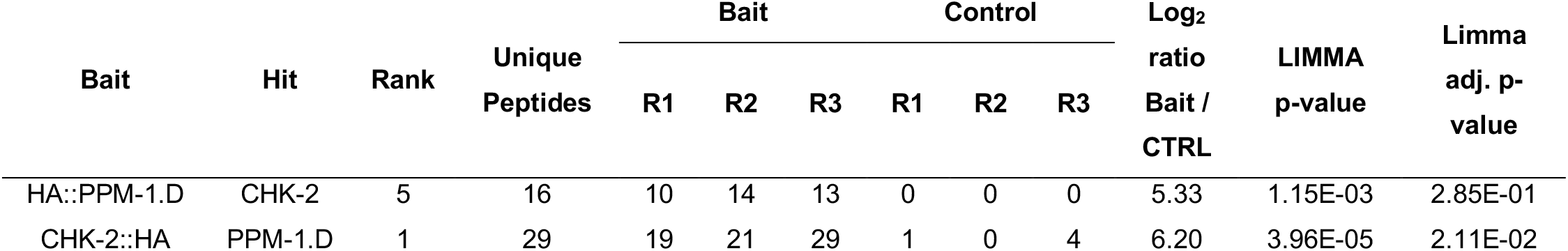
Peptide spectrum match for the bait and control indicating how often peptide of a given protein was identified in each biological replicate. Rank corresponds to the position of the identified protein when proteins are sorted by their abundance (log2 ratio bait over control). Statistical analysis was done using LIMMA T-test.

| Bait | Hit | Rank | Unique<br>Peptides | Bait |  |  | Control |  |  | Log<sub>2</sub><br>ratio<br>Bait /<br>CTRL | LIMMA<br>p-value | Limma<br>adj. p-value |
| --- | --- | --- | --- | --- | --- | --- | --- | --- | --- | --- | --- | --- |
|  |  |  |  | R1 | R2 | R3 | R1 | R2 | R3 |  |  |  |
| HA::PPM-1.D | CHK-2 | 5 | 16 | 10 | 14 | 13 | 0 | 0 | 0 | 5.33 | 1.15E-03 | 2.85E-01 |
| CHK-2::HA | PPM-1.D | 1 | 29 | 19 | 21 | 29 | 1 | 0 | 4 | 6.20 | 3.96E-05 | 2.11E-02 |

**Table S3.** P values of the Fisher’s Exact test for testing the number of RAD-51 in the mentioned mutants against the wild type. P values below 0.05 are highlighted in bold.

| Zone | Genotype | Number of RAD-51 foci |  |  |  |  |  |
| --- | --- | --- | --- | --- | --- | --- | --- |
|  |  | 0 | 1 | 2-3 | 4-6 | 7-12 | >12 |
| 1 | *ppm-1.D(tm8369)* | 0.6441 | >0.9999 | 0.2907 | >0.9999 | >0.9999 | >0.9999 |
|  | *ppm-1.D(jf120)* | 0.0589 | 0.6462 | **0.0455** | 0.1046 | >0.9999 | >0.9999 |
| 2 | *ppm-1.D(tm8369)* | **0.0014** | 0.054 | **0.006** | >0.9999 | >0.9999 | >0.9999 |
|  | *ppm-1.D(jf120)* | **0.0038** | **0.0155** | 0.3044 | >0.9999 | >0.9999 | >0.9999 |
| 3 | *ppm-1.D(tm8369)* | **0.0003** | 0.3613 | **0.0003** | **0.0259** | >0.9999 | >0.9999 |
|  | *ppm-1.D(jf120)* | 0.653 | 0.5834 | 0.6181 | **0.0168** | >0.9999 | 0.421 |
| 4 | *ppm-1.D(tm8369)* | **0.0002** | 0.0756 | 0.4791 | **<0.0001** | **0.0471** | 0.6112 |
|  | *ppm-1.D(jf120)* | **<0.0001** | **0.0006** | 0.9254 | **<0.0001** | **<0.0001** | >0.9999 |
| 5 | *ppm-1.D(tm8369)* | **<0.0001** | 0.5005 | **0.0007** | **0.002** | **0.0213** | **0.0155** |
|  | *ppm-1.D(jf120)* | **<0.0001** | 0.5292 | **<0.0001** | **<0.0001** | **0.027** | >0.9999 |
| 6 | *ppm-1.D(tm8369)* | **0.0027** | **0.0181** | >0.9999 | 0.1246 | 0.4997 | 0.2496 |
|  | *ppm-1.D(jf120)* | **<0.0001** | **0.0011** | **0.0067** | 0.0623 | >0.9999 | >0.9999 |

**Table S4.**
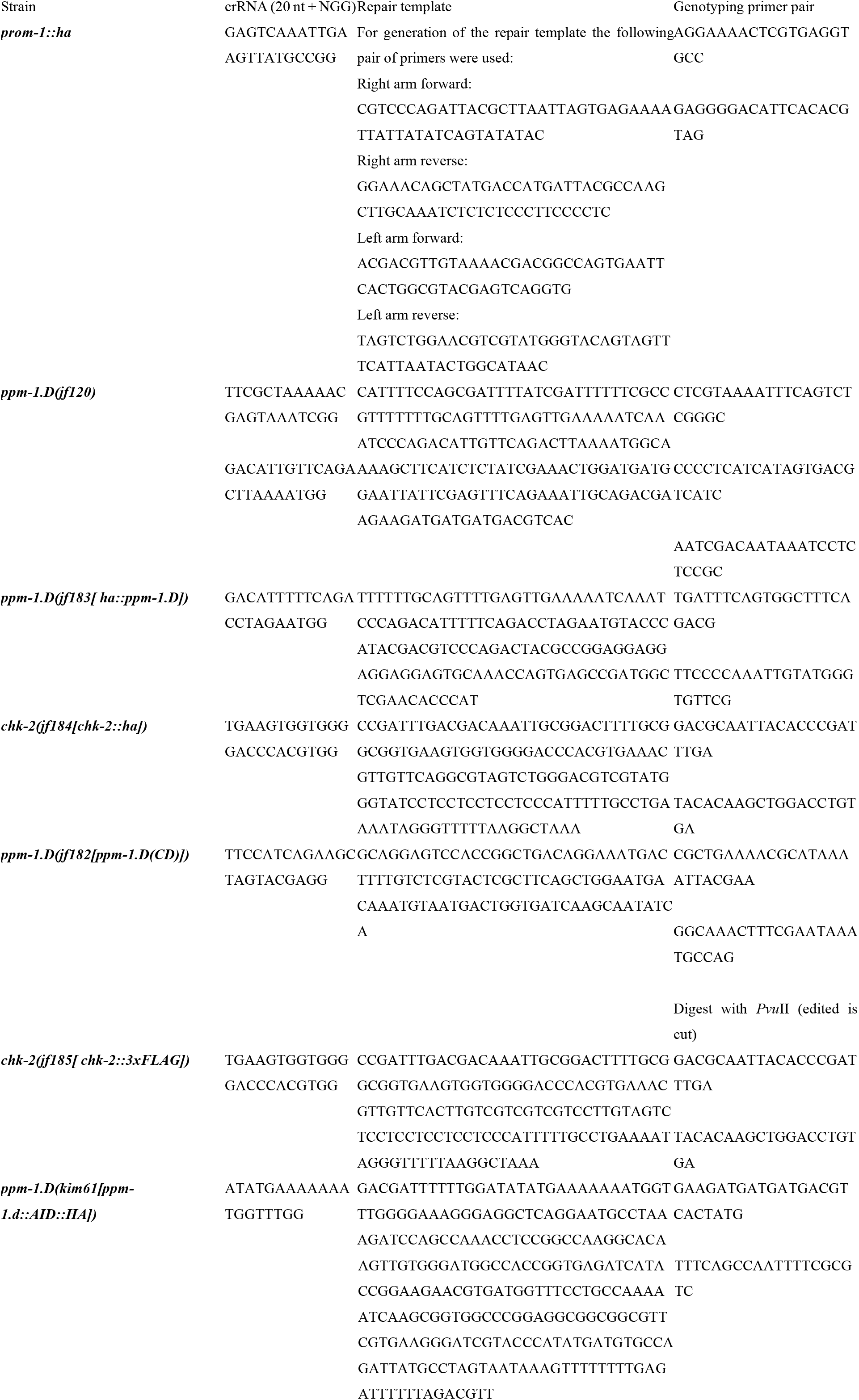
Guide, repair template and genotyping primers used in this study in 5’ to 3’ orientation as DNA sequences.

**Table S5.**
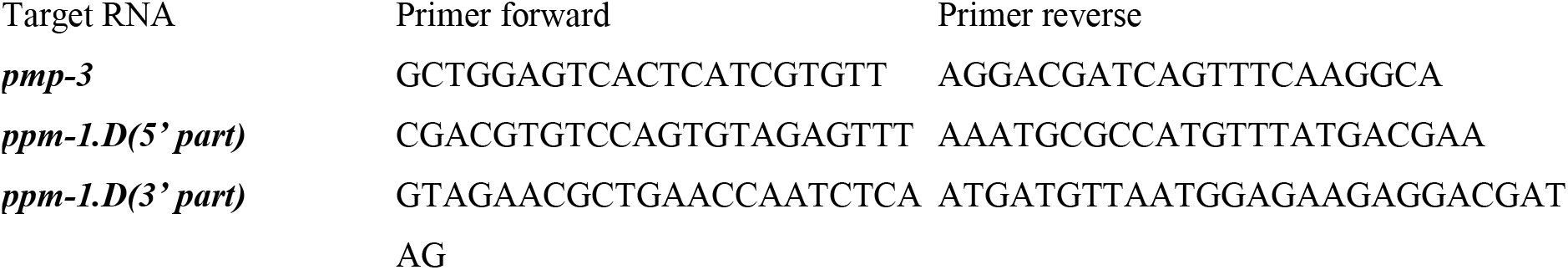
Primer pair used for qPCR.

